# Genome-wide metabolite quantitative trait loci analysis (mQTL) in red blood cells from volunteer blood donors

**DOI:** 10.1101/2022.09.07.506936

**Authors:** Amy Moore, Michael P Busch, Karolina Dziewulska, Richard O. Francis, Eldad A. Hod, James C Zimring, Angelo D’Alessandro, Grier P Page

## Abstract

The Red Blood Cell (RBC)-Omics study, part of the larger NHLBI-funded Recipient Epidemiology and Donor Evaluation Study (REDS-III), aims to understand the genetic contribution to blood donor RBC characteristics. Previous work identified donor demographic, behavioral, genetic and metabolic underpinnings to blood donation, storage, and - to a lesser extent - transfusion outcomes, but none have yet linked the genetic and metabolic bodies of work. We performed a Genome-Wide Association (GWA) analysis using RBC-Omics study participants with generated untargeted metabolomics data to identify metabolite quantitative trait loci (mQTL) in RBCs. We performed GWA analyses of 382 metabolites in 243 individuals imputed using the 1000 Genomes Project phase 3 all-ancestry reference panel. Analyses were conducted using ProbABEL and adjusted for sex, age, donation center, number of whole blood donations in the past two years, and first ten principal components of ancestry. Our results identified 423 independent genetic loci associated with 132 metabolites (p < 5×10^−8^). Potentially novel locus-metabolite associations were identified for FLVCR1 and choline, and for LPCAT3 and the lysophosphatidylserine 16.0, 18.0, 18.1, and 18.2; these associations are supported by published rare disease and mouse studies. We also confirmed previous metabolite GWA results for associations including N(6)-Methyl-L-lysine and PYROXD2, and various carnitines and SLC22A16. Association between pyruvate levels and G6PD polymorphisms was validated in an independent cohort and novel murine models of G6PD deficiency (African and Mediterranean variants). We demonstrate that it is possible to perform metabolomics-scale GWA analyses with a modest, trans-ancestry sample size.

**Key points:**

- Metabolite heterogeneity in fresh (<14 day old) RBCs donated by volunteer donors is linked to genetic polymorphisms;
- We report 2,831 high-confidence SNP-metabolite linkages (p < 5.0 × 10^−8^). Pyruvate levels in fresh RBCs are associated with glucose-6-phosphate dehydrogenase (G6PD) status

## Introduction

Red blood cells (RBCs) are the most abundant cell in the human body, representing approximately 84% of the ~30 trillion host cells in an adult individual).^1^ Despite the lack of nuclei and organelles, RBCs are endowed with around 3000 proteins^2–4^ that allow them to take up and metabolize gases (O_2_, CO_2_) and small molecule substrates. This task is facilitated by their transit through the whole body every ~20 seconds, during the average 120 day lifespan of RBCs.^5^ From this perspective, RBC metabolism has been leveraged in clinical chemistry assays as a window on systems metabolism and dysregulation thereof.^6,7^ The recent implementation of cost-effective high-throughput mass spectrometry-based metabolomics^8^ has fueled the efforts towards personalized medicine.

RBC transfusion is a life-saving intervention for 4.5 million Americans every year. The logistics of producing over 100 million units of blood available for transfusion every year around the world necessitates storage of RBC components in the blood bank. However, storage in the blood bank is characterized by a series of biochemical^9^ and morphological alterations^10^, collectively referred to as the storage lesion,^11^ which ultimately lower the efficacy of the transfusion therapy (e.g., hemoglobin increment upon transfusion).^12^ Alterations to RBC energy and redox metabolism are a hallmark of the storage lesion.^13,14^ Metabolic markers of RBC storage quality have been identified, including markers of the RBC propensity to hemolyze spontaneously^15^ or following oxidant or osmotic insults,^16^ markers of the propensity of end of storage RBCs to circulate at 24 hours upon transfusion,^17,18^ and markers of oxygen transport and off-loading function in fresh and stored RBCs.^19^ Appreciation for the role of RBC metabolism in storage quality and post-transfusion performances has informed the concept of the metabolic age of the unit – as opposed to the chronological age of the unit (i.e., days elapsed since the time of donation).^20^ Levels of metabolic markers of RBC storage quality and function are impacted by blood processing, storage additives, donor demographics (e.g., sex, race-ethnicity, age), dietary metabolites^21^ or environmental factors/donor habits (e.g., smoking, alcohol use, drugs),^22–25^ donor age^26^ and body mass index,^27^ and – relevant to the present study – donor genetics.^28^ In a move towards personalized transfusion medicine, the Recipient Epidemiology and Donor Evaluation Study – REDS III RBC Omics - was designed to test the hypothesis that donor biology plays a significant role in the quality of donated RBC.^29,30^ As part of this study, genomics approaches were used to characterize 12,353 volunteer donors enrolled at 4 different blood centers across the United States of America.^29,30^ Genetic heterogeneity of blood donors was associated with the RBC propensity to hemolyze spontaneously, or following oxidative, osmotic or mechanical insults. Quantitative trait-loci analyses were used to identify polymorphic genes that contribute to an increased resistance/susceptibility to RBC hemolysis.^28^ Of note, polymorphisms associated with a compromised (< 10%) residual activity of glucose 6-phosphate dehydrogenase (G6PD) were associated with an increased susceptibility of stored RBCs to lyse following oxidant insults.^28^ This observation was corroborated by findings from ex vivo studies,^31^ and in vivo determination of autologous post-transfusion recovery,^17^ as well as decreases in hemoglobin increments upon transfusion of G6PD-deficient units.^32^ Of note, G6PD is the rate-limiting enzyme of the pentose phosphate pathway, the main antioxidant pathway in RBCs. Thus, G6PD is crucial for the synthesis of the reducing cofactor NADPH, which is required to preserve glutathione homeostasis and reduce multiple antioxidant enzymes as they exert their catalytic activities.^33^ G6PD deficiency is common in routine blood donors of African descent (up to 13% prevalence in some metropolitan areas like New York).^34^ In G6PD deficient donors, higher levels of metabolic markers of the storage lesion have been reported, such as a faster rate of purine deamination,^35^ asparagine deamidation and methylation,^36^ and lipid oxidation.^16^ However, to date, linkage of genetic polymorphisms to metabolic heterogeneity in freshly donated blood has been limited to twin studies.^15^

Blood donors are a selected population, with the basic requirement for volunteer blood donation being the absence of serious underlying medical conditions, adequate hemoglobin levels, absence of risk factors for transfusion-transmitted infections, and not taking specific teratogenic or other medications^22^ that are grounds for deferral. As such, epidemiology studies have been facilitated by the study of large cohorts of blood donors, similar to investigations on SARS-CoV-2 incidence in the general population based on serological characterization of routine blood donors.^37^ Here we leveraged genomic^28^ and metabolomic data^29^ generated as part of the REDS III – RBC Omics study to perform a metabolite Quantitative Trait Loci (mQTL) analysis of routine blood donors. The study builds on previous mQTL reports in the context of cardiovascular diseases, asthma, or neurodegenerative diseases, including Alzheimer’s disease.^38–42^ Given the importance of RBC metabolism as a window into systems homeostasis, findings reported here could be relevant not just for transfusion medicine research, but also for diverse areas of physiology where RBC metabolism is impaired (e.g., exercise,^43^ aging,^44^ adaptation to high-altitude hypoxia,^45^ pathological hypoxia upon hemorrhage,^46^ COVID-19,^47,48^ cardiovascular^49^ and kidney diseases^50,51^).

## Methods

### REDS-III RBC-Omics study participants and samples

RBC-Omics was conducted under regulations applicable to all human subject research supported by federal agencies as well as requirements for blood product manipulation specified and approved by the FDA. The data coordinating center (RTI International) of REDS-III was responsible for the overall compliance of human subjects regulatory protocols including institutional review board approval from each participating blood center, from the REDS-III Central Laboratory (Vitalant Research Institute), and from the data coordinating center, as previously detailed.^16,29^ Donors were enrolled at the four participating REDS-III US blood centers. Overall, 13,758 whole blood donors were enrolled and 13,403 (97%) age 18+ provided informed consent to participate in the study; of these, 12,353 were evaluated for hemolysis parameters (spontaneous, oxidative or osmotic) on RBCs stored for ~39-42 days. Extreme hemolyzers (5^th^ and 95^th^ percentile) from the donors tested for end of storage oxidative hemolysis were asked to donate a second unit of blood. These units were sterilely sampled for metabolomics analyses (n = 250 for the freshest available time points, i.e., < 14 storage days). Blood collection, sample processing and other aspects of the screening and recall phases of the RBC-Omics Study have been extensively described.^26,28^

### Sample processing and metabolite extraction

An isotopically labeled internal standard mixture including a mix of ^13^C^15^N-labeled amino acid standards (2.5 μM) was prepared in methanol. A volume of 100μl of frozen RBC aliquots was mixed with water and the mixture of isotopically labeled internal standards (1:1:1, *v/v/v*). The samples were extracted with methanol (final concentration of 80% methanol). After incubation at −20°C for 1 hour, the supernatants were separated by centrifugation and stored at −80°C until analysis. Samples were vortexed and insoluble material pelleted as described.^16,29^

### Ultra-High-Pressure Liquid Chromatography-Mass Spectrometry metabolomics

Analyses were performed using a Vanquish UHPLC coupled online to a Q Exactive mass spectrometer (Thermo Fisher, Bremen, Germany). Samples were analyzed using a 3 minute isocratic condition or a 5, 9 and 17 min gradient as described.^52–54^ Solvents were supplemented with 0.1% formic acid for positive mode runs and 1 mM ammonium acetate for negative mode runs. MS acquisition, data analysis and elaboration was performed as described.^52–54^ Additional analyses, including untargeted analyses and Fish score calculation via MS/MS, were calculated against the ChemSpider database with Compound Discoverer 2.0 (Thermo Fisher, Bremen, Germany).

### Metabolite QC and Processing

The quality control and processing of metabolites is detailed in **Supplementary Figure 1**. We first selected only those metabolites measured at Day 10 of storage, for which 250 participants had metabolite data. Metabolites with missing data and zeros were both treated identically as missing. We removed the following metabolites from further processing: a duplicate carnosine, a duplicate lorazepam, phosphate, triacanthine, and acetyl-L-carnitine. 507 metabolites remained for further processing. We also removed 22 drug metabolites with concentrations detected in greater than 50% of the participants, leaving 487 metabolites. We then separated the participant data by blood storage additive type and excluded metabolites with greater than 10% missingness from each additive set, respectively. After removing these metabolites, 382 remained. We separated these 382 metabolites into those quantified absolutely (against stable isotope-labeled internal standards, as described^55^) vs relatively. Relatively quantified metabolites were natural log-transformed. A suffix “(uM)” was added to the label of all the metabolites for which absolute concentrations were determined. These groups of metabolites then had missing metabolite levels imputed using QRILC,^56^ implemented in the R package QRILC.^56^ After imputation, all metabolites were inverse-normal transformed using the R package GenABEL rntransform command.^57^

### Genotyping and Imputation

Details of the genotyping and imputation of the RBC Omics study participants have been previously described by Page, et al.^28^ Briefly, genotyping was performed using a Transfusion Medicine microarray^30^ and the data are available in dbGAP accession number phs001955.v1.p1. Imputation was performed using 811,782 SNPs that passed quality control. After phasing using Shape-IT,^58^ imputation was performed using Impute2^59^ with the 1000 Genomes Project phase 3^59^ all-ancestry reference haplotypes. We used the R package SNPRelate^60^ to calculate principal components (PCs) of ancestry.

### Genome-Wide Association Study

We performed association analyses for each of the 382 metabolites using an additive SNP model in the R package ProbABEL^61^ and 243 study participants who had both metabolomics data and imputation data on serial samples from stored RBC components that passed respective quality control procedures. We adjusted for sex, age (continuous), frequency of blood donation in the last two years (continuous), blood donor center, and ten ancestry PCs. Statistical significance was determined using a p-value threshold of 5×10^−8^. We only considered variants with a minimum minor allele frequency of 1% and a minimum imputation quality score of 0.80.

### Replication and Sensitivity Analyses

For replication, we followed the same procedures for post-processing of metabolites measured at Day 42 of storage. There were 176 participants with metabolite data generated from Day 42 samples. Association analyses and statistical significance were determined as described above. We selected 46 metabolites, oversampled for top hits from the GWAS analysis of early storage samples to analyze for potential replication and in the sensitivity analyses described below.

We performed four sensitivity analyses using 46 metabolites and the original 243 recalled RBC-Omics participants. We performed a “stringent” GWAS, which required that evaluated variants have a minimum minor allele frequency of 5% and a minimum imputation quality score of 0.90. We performed an analysis using only those participants whose blood donations were collected at one of the three centers that used Additive 3 in their storage protocol. We also performed a sensitivity analysis including only those participants of European ancestry and using the variant data imputed using the European reference panel. Finally, we re-imputed the missing metabolites data as described above, swapping out the QRILC imputation procedure for a simple substitution of the missing value with the lowest detected value for the metabolite in question.

### OASIS Queries

The OASIS: Omics Analysis, Search & Information a TOPMED funded resources^62^, was used to annotate the top SNPS. OASIS annotation includes information on position, chromosome, alellele frequencies, closest gene, type of variant, position relative to closest gene model, if predicted to functionally consequential, tissues specific gene expression, and other information.

### Locuszoom Plots

We generated LocusZoom plots locally using v1.4 and plotted a margin of +/− 200 kilobases around each lead SNP against the November 2014 1000 Genomes European ancestry build

### Comparison with GWAS Catalog

Lead SNPs for all metabolites were queried using the LDLink tool LDtrait (query date 5/6/2022)^63^ by selecting an R^2^ threshold of 0.8 in a +/− 500,000 base pair window in the combined five European ancestries using Genome Build GRCh37. We noted SNPs that have been previously associated with other traits and considered replicated associations as those SNPs with previously reported associations to the same metabolite as found in our study population.

### Animal models

Humanized G6PD deficient (A-, Med-) and non-deficient (HuCan) mice were generated by replacing the murine G6PD locus in Bruce4 ES cells (C57BL/6 background) with either the A-(V68M/N126D), Med- (S188F) or huCan (B+) variant (manuscript in preparation). In short, nucleofected ES cells were drug-selected (Neo), G418 resistant clones were isolated, and the presence of homologous recombination (and absence of random integration) was confirmed (data not shown). Clones were then developed into full animals and correct homologous recombination was reconfirmed. The Neo cassette was flanked with FRT sites and removed by breeding with a germline FLP transgenic mouse – the FLP was subsequently removed. Generation of Cre-inducible G6PD Med-deficient mice was described previously. ^64^

### G6PD deficient subjects

Male volunteers were recruited using flyers, person-to-person communication, and email, between November 2012 and August 2017. Screening was limited to males because G6PD deficiency is X-linked; Following screening and confirming G6PD activity, 10 G6PD-deficient and 30 G6PD-normal males donated 1 unit of whole blood at the New York Blood Center (New York, New York), ^17^ each of which was processed into packed RBCs, leukoreduced, prior to metabolomics analysis.

## Results

Metabolomics analyses were performed on packed RBC samples derived from stored RBC components from 250 donors who had been previously characterized at the genome level via the Precision Transfusion Medicine array **(Figure 1.A)**.^30^ Through the workflow summarized in **Supplementary Figure 1,** a total of 2,831 SNP-metabolite associations were observed below the genome-wide correction threshold (p < 5.0 × 10^−8^). Data are summarized in tabulated form in **Table 1** by identifying the SNP with the smallest p-value within a +/− 500 kilobase range as the lead SNP; individual SNP associations are reported extensively in **Supplementary Table 1**. In **Figure 1.B**, we listed the top 10 hits identified by closest annotated gene to the significant SNP in order of - log10(p). Manhattan plots overlapping all the significant hits (FDR < 5 × 10^−8^) are shown in **Figure 1.C**, which also includes metabolite – gene pairs. QQplots for the top nine metabolite-associated SNPs are shown in **Figure 1.D**. Sensitivity analyses for 46 metabolites examining the impact of 1) more stringent variant QC; 2) the choice of imputation strategy for missing metabolite data; 3) the effect of blood storage additive; and 4) ancestry, are reported in **Supplementary Table 2**. Genetic associations identified for six metabolites have been previously reported (**Supplementary Table 3)**. Ancestry plots were thus generated to show normalized metabolite abundances as a function of alleles, as distributed across genetic ancestries of the donors enrolled in this study (**Figure 2.A**). We further characterized the mQTL loci by generating LocusZoom plots to examine the local LD structure and performed *in silico* functional annotation using OASIS.

**Figure 1.**
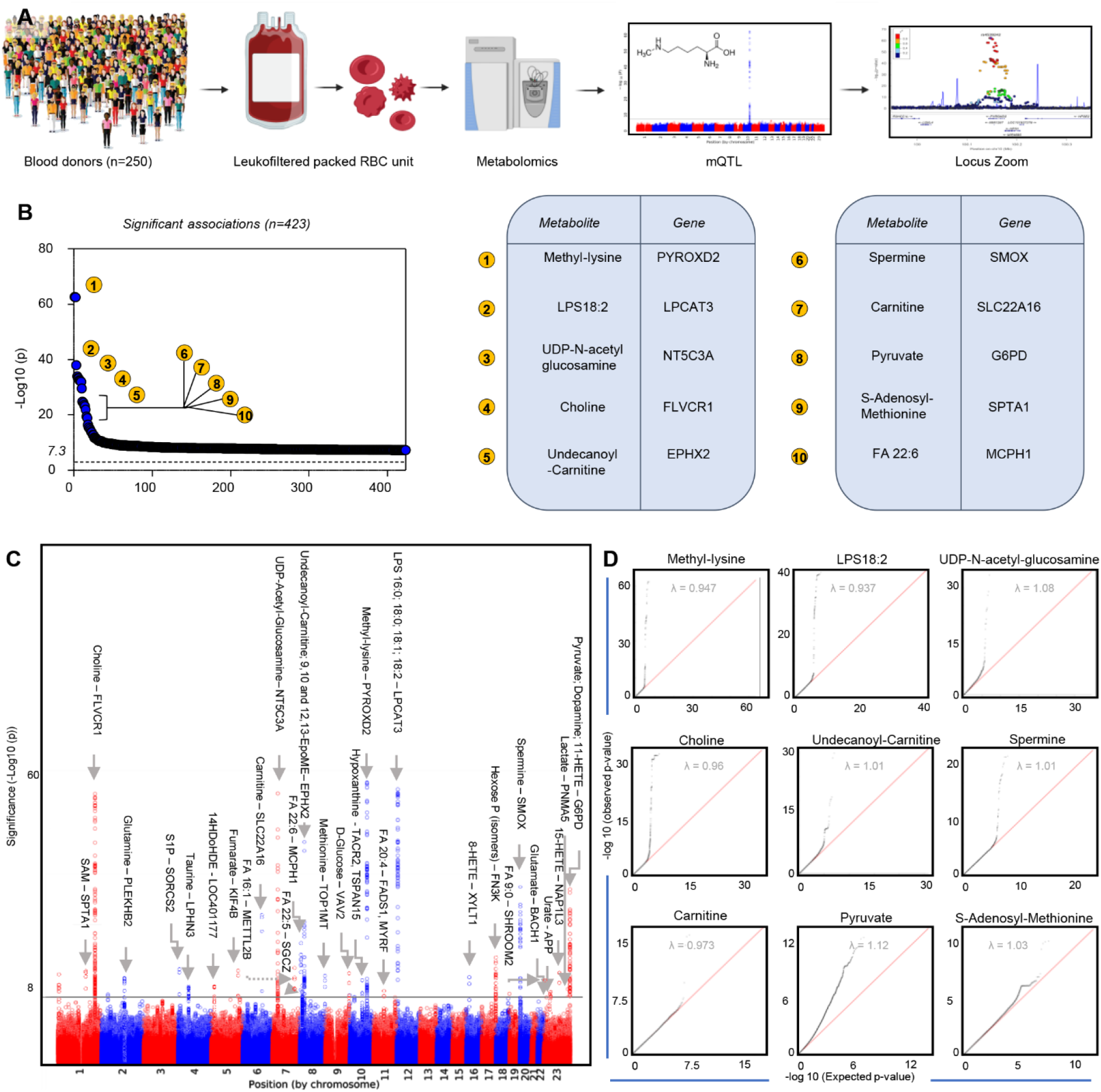
Study design and top 10 hits from the mQTL analysis from the REDS III RBC omics pilot recalled donor study. Metabolomics analyses were performed on 250 packed RBC samples from donors who had been previously characterized at the genome level via the Precision Transfusion Medicine array **(A)**.^30^ In **B**, an overview of the top 10 hits (closest annotated gene to the identified SNP) as a function of significance (-log10(p)). In **C**, overlapped Manhattan plots of all the significant hits (FDR < 5 × 10^−8^), including metabolites – gene pairs. Each data point corresponds to a -log10(P value) from a multivariant linear regression model’s P value for an SNP. The black horizontal line represents an accepted P-value level of genome-wide significance (P = 5 × 10^−8^). In **D**, qqplots for the top 10 hits from the mQTL analysis.

**Figure 2.**
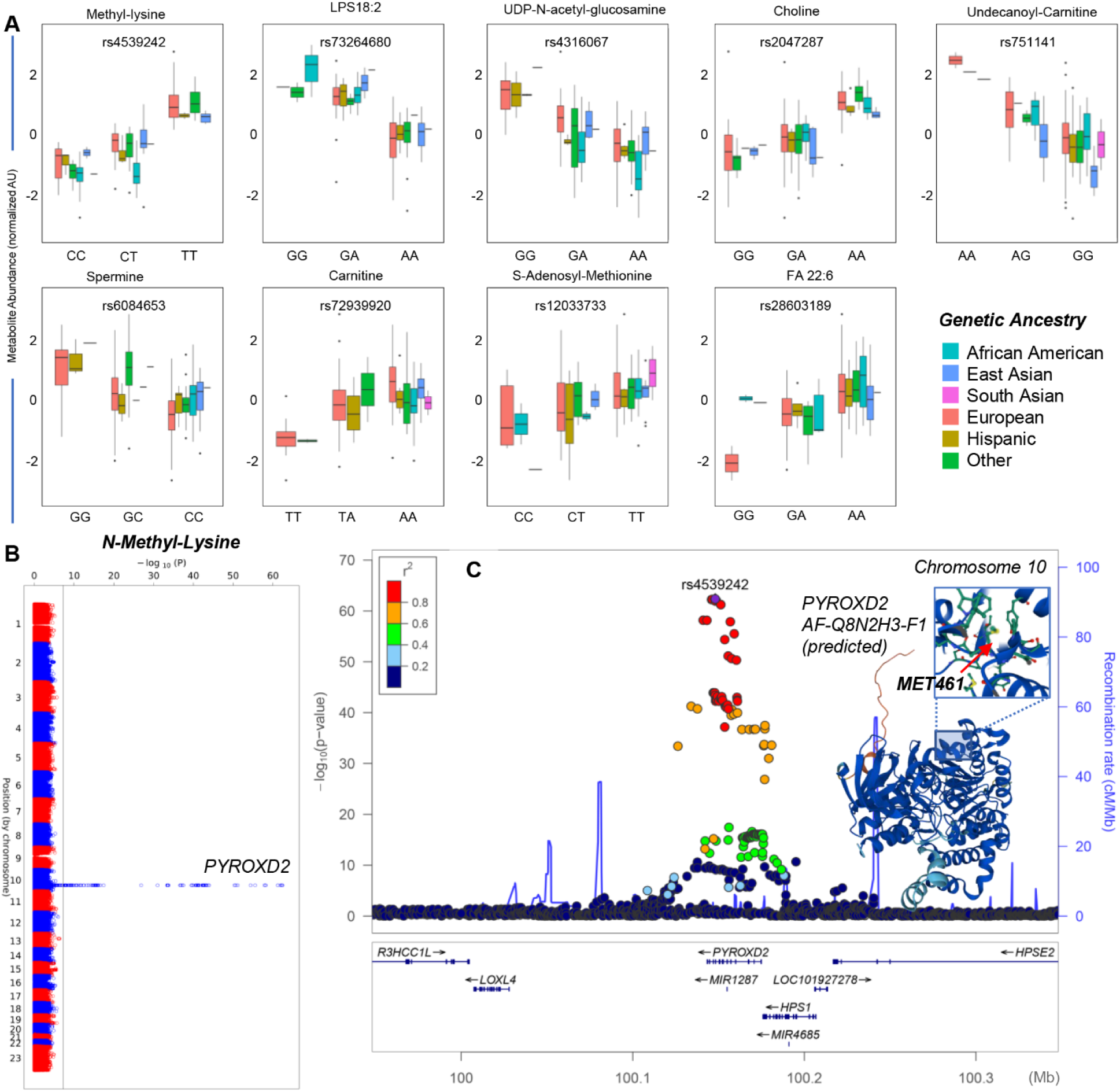
Ancestry plots and association between methyl-lysine levels and polymorphisms in PYROXD2. In **A**, for the top GWA hits we generated box and whisker plots based on metabolite abundances as a function of allele variance across all genetic ancestries in this study. Consistently with previous mQTL studies,^40,41,65–67^ polymorphisms in the exonic region coding for the enzyme PYROXD2 were associated with variance in the levels of methyl-lysine, an observation that represents a sort of internal quality control for the present analysis compared to the literature. Manhattan plots and LocusZoom are shown in panels **B-C**, respectively.

Consistent with previous mQTL studies(citations 40,41,64–67), the top SNP associated with levels of methyl-lysine, rs4539242, is in high linkage disequilibrium (LD) with both the missense mutation M461T (R^2^=1.0 in Europeans) and synonymous mutation F484F (R^2^=0.94 in Europeans), observations that represent an internal quality control for the present analysis (Manhattan plots and LocusZoom in **Figure 2.B-C**, respectively). Both mutations are themselves associated with levels of methyl-lysine (p=4.22×10^−13^ and p=1.28×10^−44^, respectively; **Supplementary Table 1**).

The region coding for the enzyme lysophosphatidylserine acetyl-transferase 5 (LPCAT3) was found to be genetically heterogenous across volunteer blood donors. Polymorphisms in the region coding for LPCAT3 were The lead SNP, rs73264680, associated with associated with RBC levels of lysophospholipids (LPS), including linoleyl- (18:2), palmitoyl (16:0), stearoyl (18:0) or oleyl-LPS (18:1), rs73264680, is in perfect LD in Europeans with the missense mutation rs1984564/I217T within *LPCAT3* (**Figure 3.A-C; Supplementary Table 2**; residue mapped against the structure of LPCAT3 - 7F3X.pdb in **Figure 3.D**).

**Figure 3.**
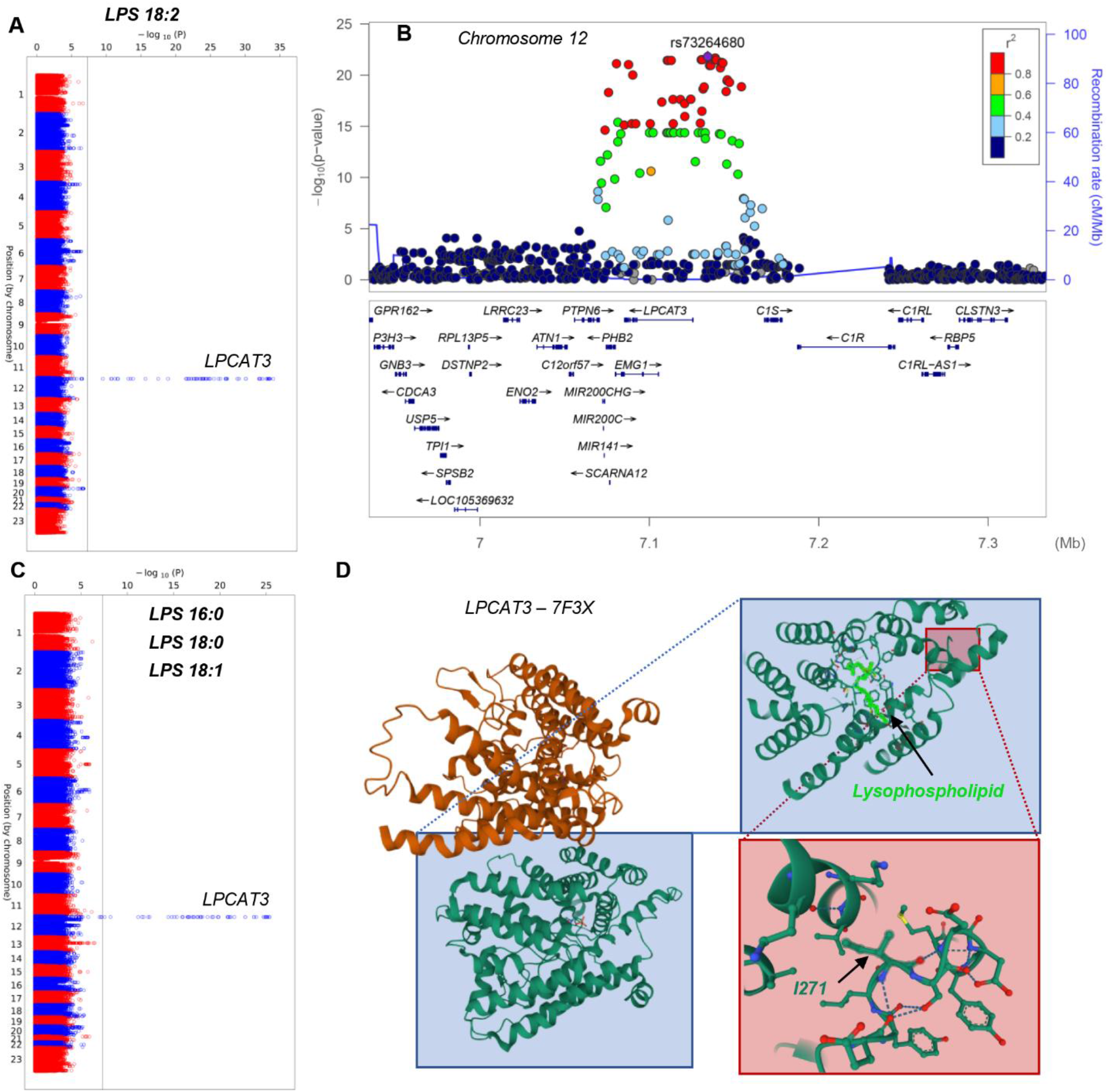
LPCAT3 is polymorphic in healthy blood donors and associates with red blood cell lysophospholipid (LPS) levels. Manhattan plots for LPS, specifically linoleyl-(18:2 - **A** and related LocusZoom, highlighting the association with the region coding for LPCAT3 in **B**), palmitoyl (16:0), stearoyl (18:0) or oleyl (18:1 - **C**). In **D**, highlight of the polymorphic residue I271, mapped against the structure of LPCAT3 (7F3X.pdb).

The lead SNP associated with UDP N acetyl glucosamine, rs4316067 (Table 1), is located in an intron of *NT5C3A* (**Figure 4A-B**). The lead SNP associated with choline, rs2047287 (**Figure 4C-D**), is in strong LD (D’=1.0; R2=0.768 in Europeans) with the missense mutation T544M in *FLVCR1*.

**Figure 4.**
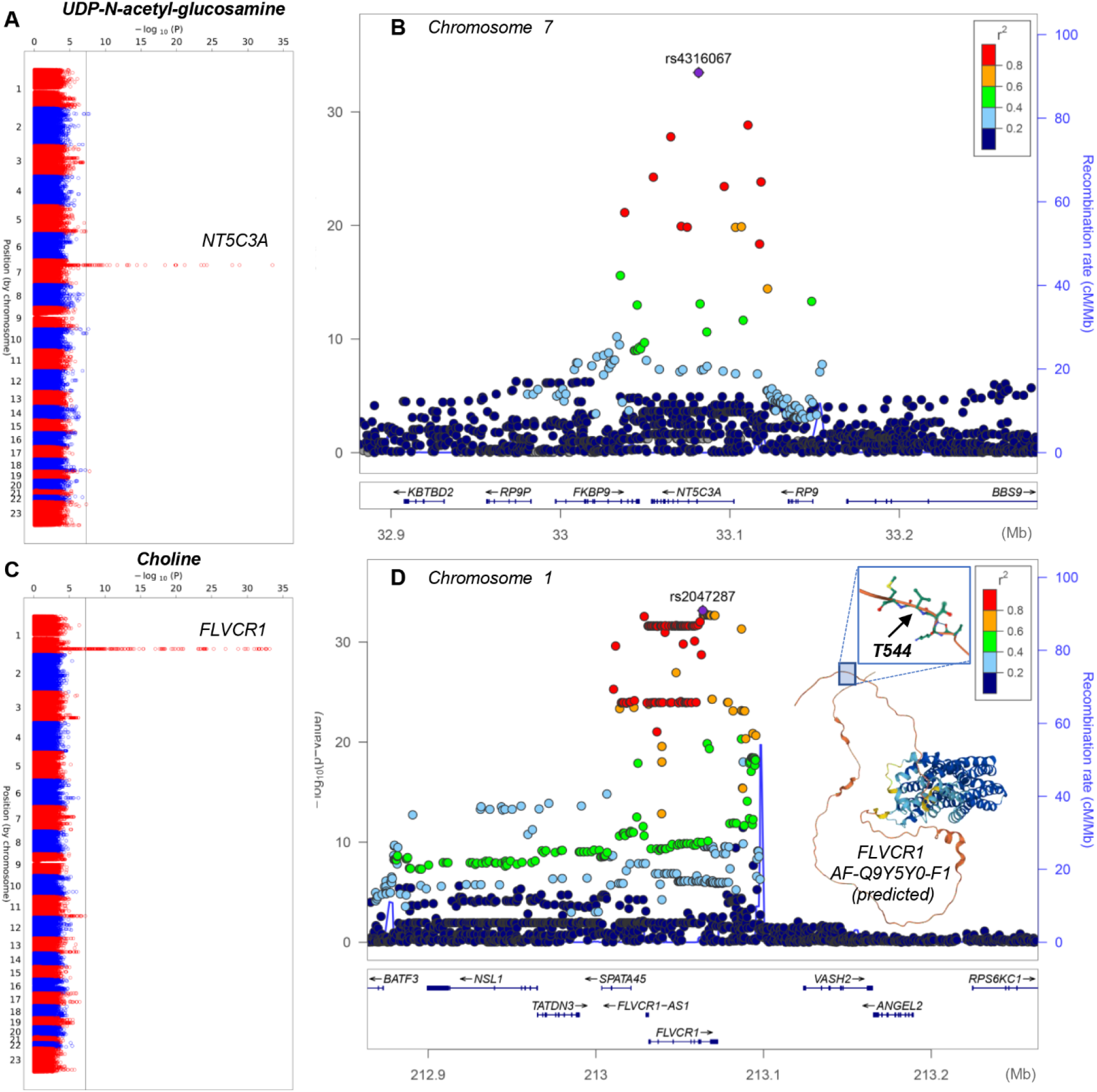
Polymorphisms in NT5C3A and FLVCR1 are associated with variability in the levels of UDP-N-acetyl-glucosamine and choline in RBCs from healthy blood donors. Manhattan plots and LocusZooms are showns in panels **A-B** and **C-D**, respectively.

Polymorphisms in bifunctional epoxide hydrolase 2 (*EPHX2* - missense mutation rs751141/R221Q - p = 4.55 10^−10^; λ = 1.01) and spermine oxidase, where the intronic rs11087622 in *SMOX* – is in LD (R^2^=0.16; D’=0.71 in Europeans) with synonymous mutation A392A (**Supplementary Table 1**)-, are associated with variability in the levels of oxylipins (12,13-EpOME) and spermine, respectively (**Figure 5**).

**Figure 5.**
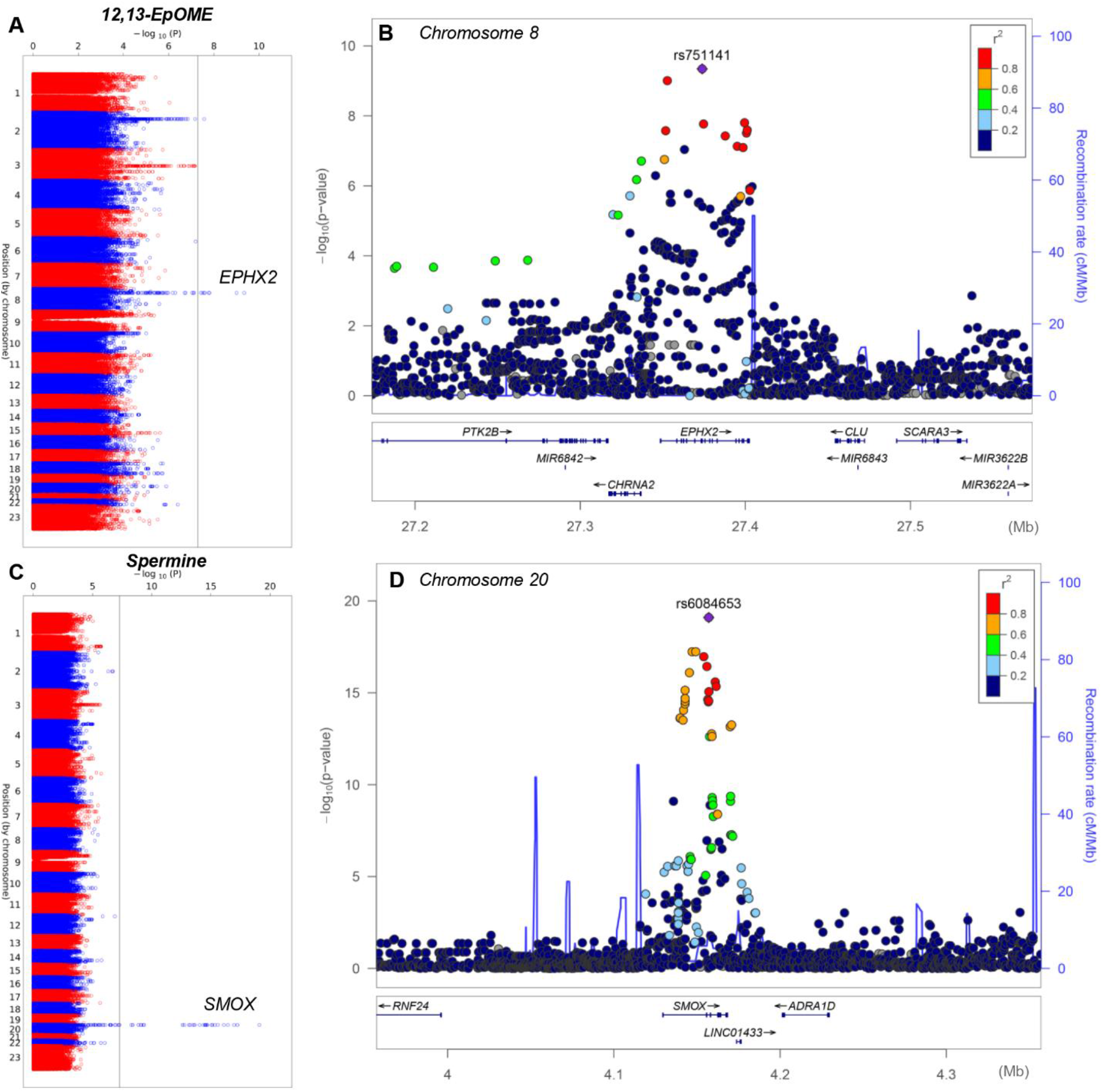
Polymorphisms in EPHX2 and SMOX are associated with variability in the levels of oxylipins (12,13-EpOME) and spermine, respectively. Manhattan plots and LocusZooms are showns in panels **A-B** and **C-D**, respectively.

A missense variant (rs12210538/M409T) in the carnitine transporter *SLC22A16* is associated with variability in the levels of RBC free and acetyl-carnitine (**Supplementary Table 1**). Manhattan plots and LocusZooms are shown in **Supplementary Figure 2.** Additional SNPs associated with carnitine levels include palmitoyl-carnitine (nearest gene *HTR5A-AS1*) and undecanoyl-carnitine (nearest gene *EPHX2*).

A series of significant associations were identified between the levels of oxylipins like 9.10-EpOME (EPHX2 – Uniprot names provided for protein products of the nearest gene within parentheses in this paragraph) or 14-DHoHE (LOC401177 or LOC100505817) or 9-HETE, 15-HETE (NAP1L3), dopamine (G6PD), glycolytic metabolites (glucose and VAV2, hexose phosphate, including fructose 6-phosphate and FN3K; lactate and PNMA5), purines (hypoxanthine and TACR2, TSPAN15; urate and FOLR1 and APP), amino acids (glutamine and PLEKHB2; glutamate and BACH1-IT2; methionine and TOP1MT; taurine and LPHN3, threonine and mR8058), free fatty acids (palmitoleic acid and METTL2B; oleic acid; arachidonic acid and FADS1; docosapentaenoic acid and SGCZ), sphingolipids (sphingosine 1-phosphate and EDARADD or SORCS2 or KDM6A), uridine diphosphate (UDP and ZNF485 – **Supplementary Figure 3-15**).

Finally, polymorphisms in *SPTA1* and *G6PD* are associated with variability in the levels of S-adenosyl-methionine (intronic - p = 2.52 10^−10^; λ = 1.03; **Supplementary Table 1**) and pyruvate **μM** (missense mutation V98M - p = 2.87 10^−12^; λ = 1.12; **Supplementary Table 1**), respectively (**Figure 6.A-B** and **C-D** for Manhattan plots and LocusZoom).

**Figure 6.**
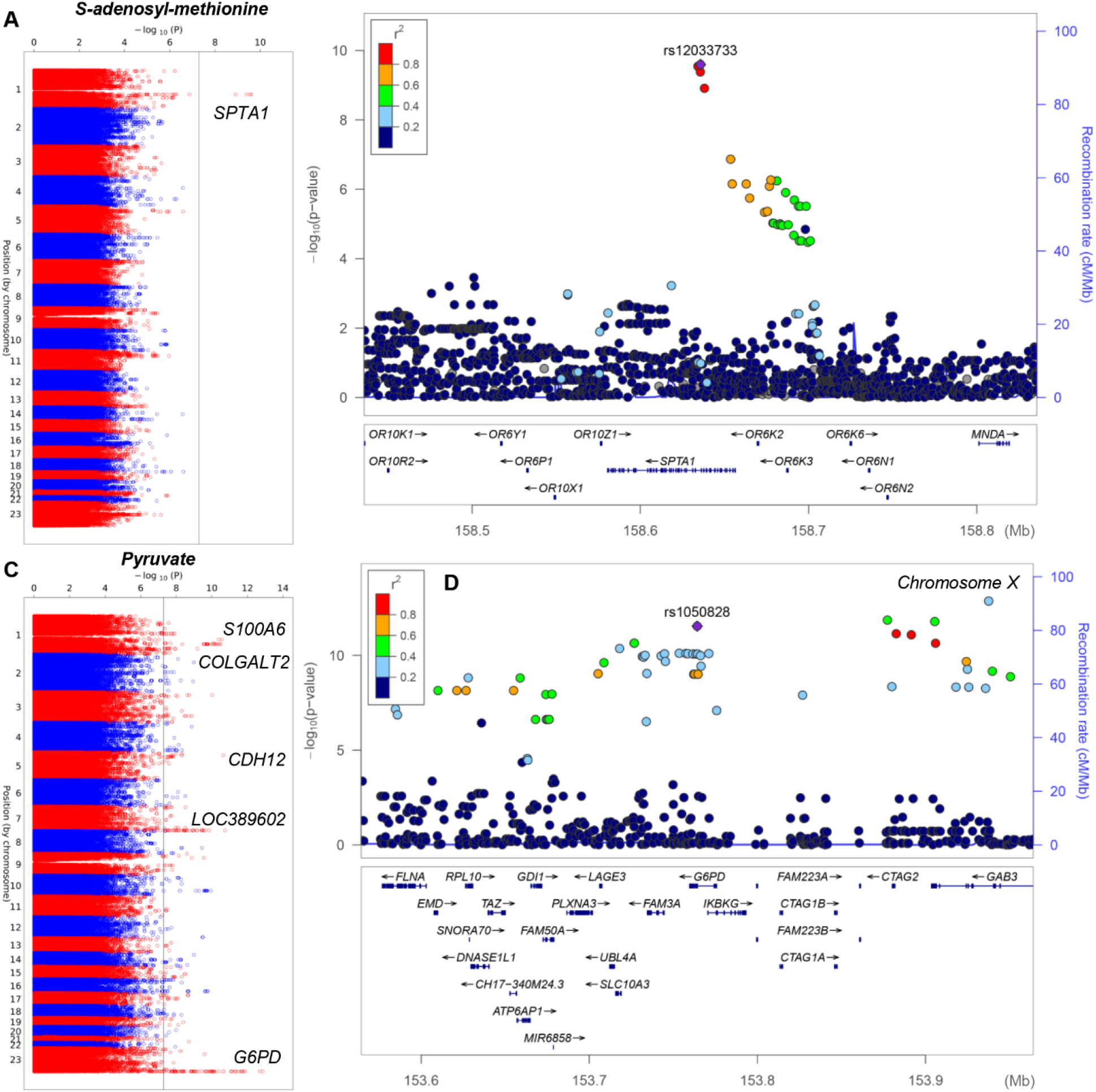
Polymorphisms in SPTA1 and G6PD are associated with variability in the levels of S-adenosyl-methionine and pyruvate, respectively. Manhattan plots and LocusZooms are showns in panels **A-B** and **C-D**, respectively.

### Sensitivity analyses

We performed several sensitivity analyses, including GWAS of 46 metabolites in 176 study participants with available metabolomics data generated from RBC samples stored for 42 days. The top associations for N6-Methyl-L-Lysine, LPS16.0-18.2, UDP N-aceytl-glucosamine, Choline, Undecanoyl carnitine, spermine, spermine uM, and L-carnitine, replicated in each of the five sensitivity analysis, with either the original lead SNP or a genome-wide significant SNP in high LD with the original reaching genome-wide significance (**Supplementary Table 1**). For docosahexaenoic acid (FA22.6), the association with rs28603189 replicated when stringent QC criteria were applied, when a different missing metabolite imputation strategy was employed, and when analysis was restricted to the participants whose RBC samples were stored in Additive 3 (R^2^=0.94), but not in the day 42 storage samples or the European-only analysis. The association between rs12033733 and S-adenosyl-L-methionine withstood the stringent QC but none of the other sensitivity analyses. Finally, although there were many genome-wide significant associations with pyruvate uM, the lambda was 1.115 and the QQ plot troubling (**Figure 1**), potentially indicating unaccounted for population structure for this metabolite. The association between pyruvate uM and rs142516556, a SNP near the *G6PD* gene, remained robust to the stringent QC, imputation method, and Additive 3 restricted GWAS (**Supplementary Table1**).

### Elevated pyruvate and pyruvate/lactate ratios are recapitulated in an independent human cohort of blood donors and mouse models of G6PD deficiency

Pyruvate levels were found to be inversely proportional to G6PD activity in fresh RBCs from an independent cohort of G6PD deficient (n=10) and sufficient (n=27) blood donors (**Figure 7.A-C**). The differences in pyruvate levels between the two groups were exacerbated during blood bank storage up to 42 days (**Figure 7.D**). Similarly, RBCs from G6PD deficient mice (African A- and Mediterranean variant – Med-) and WT C57BL6/J or humanized canonical G6PD mice (**Figure 7.E**) were incubated with 1,2,3-^13^C_3_-glucose for 1h, to determine metabolic fluxes through glycolysis and the pentose phosphate pathway (PPP). Results (**Figure 7.F**) confirmed significant decreases in the labeled levels of oxidative phase metabolites of the PPP (^13^C_3_-phosphogluconate and ^13^C_2_-ribose-phosphate) in A- and Med-mice, which corresponded to increases in the ratios of labeled ^13^C_3_-pyruvate/lactate.

**Figure 7.**
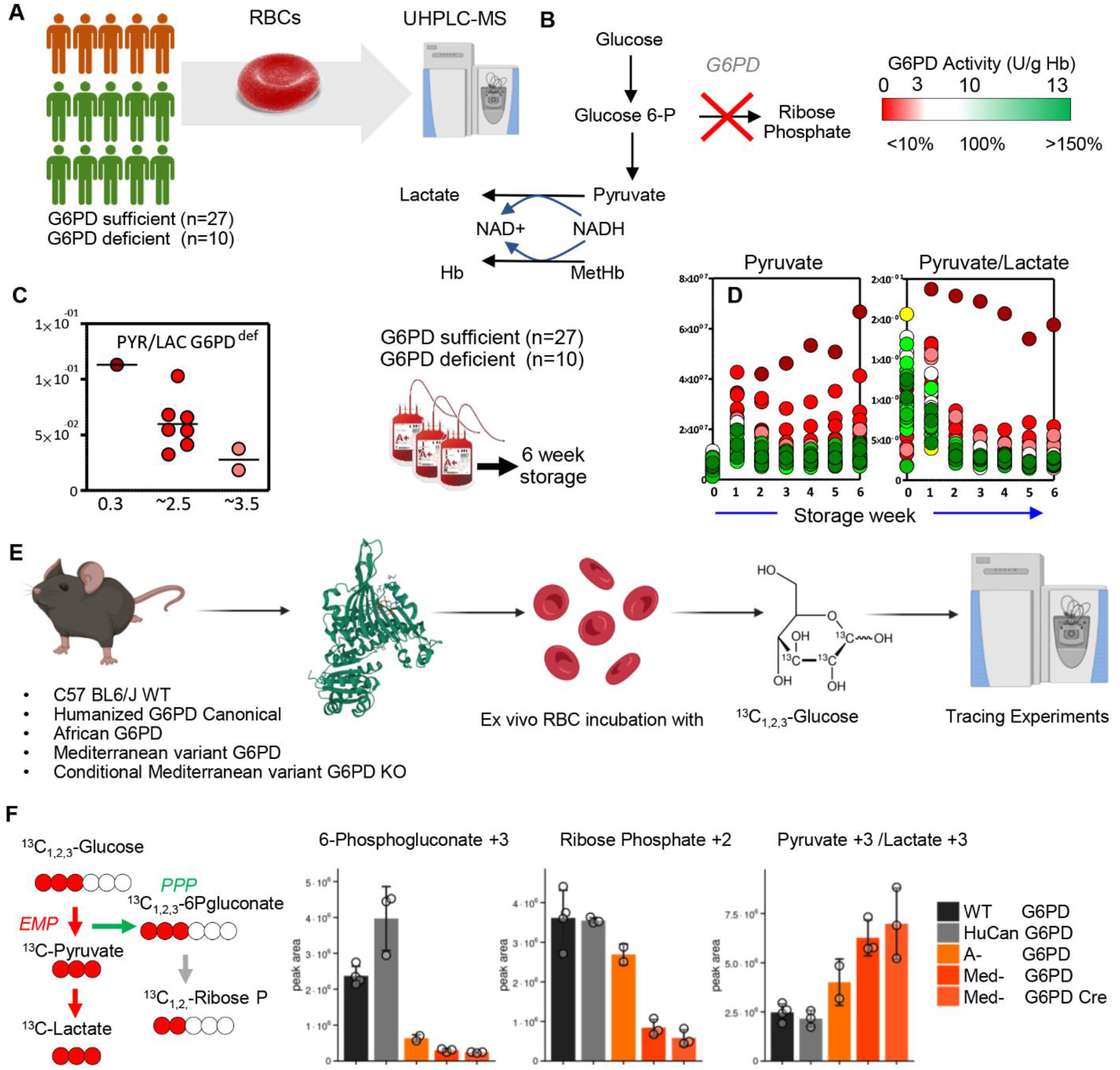
G6PD deficiency in fresh and stored RBCs from blood donors are associated with increases in pyruvate levels and pyruvate/lactate ratios. Pyruvate levels were found to be inversely proportional to G6PD activity in fresh RBCs from G6PD deficient (n=10) and sufficient (n=27) blood donors (**A-C**). These differences in pyruvate levels between the two groups were exacerbated during storage in the blood bank up to 42 days (**D**). Similarly, RBCs from G6PD deficient mice (African and Mediterranean variant) and WT C57BL6/J or humanized canonical G6PD mice (**E**) were incubated with 1,2,3-^13^C_3_-glucose for 1h, to determine metabolic fluxes through glycolysis and the pentose phosphate pathway (PPP). Results (**F**) confirmed significant decreases in the labeled levels of oxidative phase metabolites of the PPP (^13^C_3_-phosphogluconate and ^13^C_2_-ribose-phosphate) in A- and Med-mice, which corresponded to increases in the ratios of labeled ^13^C_3_-pyruvate/lactate.

## Discussion

As part of the REDS-III RBC-Omics study, a cohort of 12,535 volunteer blood donors were enrolled to donate a unit of blood that was processed into RBC components that were characterized for storage hemolysis parameters. DNA samples derived from donation WBC were genotyped using a Precision Transfusion Medicine array mapping 879,000 SNPs.^30^ Phenotypes that were associated with these polymorphisms related to RBC propensity to hemolyze, either spontaneously or following oxidant or osmotic instults.^28^ As a result, 27 loci were associated with measures of hemolysis following blood storage, the most significant being the association between ANK and SPTA1 with osmotic hemolysis (5.85 × 10^−28^ and 1.01 × 10^−22^, respectively) and G6PD with oxidative hemolysis (2.66 × 10^−17^).^28^ Here we performed the first mQTL analysis of RBCs from 250 recalled RBC-Omics donors, a subset of the 12,535 enrolled and genotyped donors in the RBC-Omics study.

Overall, we report 2,831 SNP-metabolite associations meeting genome-wide significance. Of note, the smallest p-value found in the present study, p=1.90×10−63 for the association between rs4539242 within *PYROXD2* and the RBC levels of N-methyl-lysine, was much smaller than the p-values describing any of the 27 loci associated with RBC hemolysis phenotypes in this population (citation 28), despite the much smaller cohort (250 vs 12,535) – suggesting the metabolic signatures are more directly determined by genetics than is hemolytic propensity, with the latter having a larger etiologic contribution from environmental factors. The association between *PYROXD2* and methyl-lysine had already been reported in previous mQTL studies,^40,41,65–68^ and thus serves as an internal control for the present analysis.

Novel findings include the association between polymorphisms in the heme transporter FLVCR1 and the RBC levels of choline. Previous genomics data have shown a strong linkage (co-dependency: 0.41 Pearson) between FLVCR1 and the enzyme choline kinase A (https://depmap.org/portal/gene/CHKA?tab=overview). By controlling intracellular heme pools, FLVCR1 is known to play a role in the differentiation of committed erythroid progenitors.^69^ Like methionine, choline is a methyl-group donor to recharge S-Adenosyl-methionine (SAM), and thus it indirectly participates in nucleotide synthesis during erythropoiesis^70^ and methylation events to repair isoaspartyl-damage upon oxidant insults in vivo and in vitro.^71^ In this view, it is interesting to note that SAM levels were associated with polymorphisms in the structural protein spectrin alpha 1 (SPTA1), recently identified as one of the main targets of methylation of deamidated asparagines in stored RBCS.^72^ The association between choline and FLVCR1 is also relevant owing to the role of choline metabolism as a substrate for phosphocholine metabolism in phospholipid synthesis in terminal erythropoiesis.^73^

The RBC levels of multiple lysophosphatidylserines (LPS 16:0, 18:0, 18:1 and 18:2) were associated with variation in the lysophosphatidylcholine acetyl-transferase 5 gene (LPCAT3). LPCAT3 is a key enzyme of the Lands cycle,^74^ which participates in the repair of oxidatively damaged lipids, including LPS, and the secretion of triglycerides through regulation of arachidonoyl-phospholipid metabolism in RBCs.^75^ These results are suggestive of a role of LPCAT3 in phosphatidylserine metabolism, a class of lipids whose exposure in the outer membrane leaflet regulates erythrophagocytosis and clearance from the bloodstream, with implications for posttransfusion recovery of stored RBCs.^10^ In this view, it is worth noting that the levels of multiple acyl-carnitines, in equilibrium with the acyl-CoAs as part of the Lands cycle, were found to be associated with polymorphisms in the carnitine transporter SLC22A16. This observation may indicate a inter-subject variability in membrane lipid damage-repair capacity, with implications for exercise physiology or kidney disease, since this pathway is impacted by acute exercise^43^ or (hypoxia-induced) kidney dysfunction,^51^ and carnitine-containing supplements. It is worth noting that polymorphic Mitochondrial topoisomerase I (TOP1MT), which limits doxorubicin-induced cardiotoxicity^76^ and kidney dysfunction,^77^ was previously associated with inter-donor variability in the levels of methionine – suggestive of a potential axis in RBC oxidant stress damage-repair mechanisms for proteins^71^ and lipids^77,78^. Similarly, this observation may be relevant to storage quality and the implementation of carnitine-containing blood storage additives.^79^ Heterogeneity in the RBC levels of some (bacteria or, under sterile ex vivo conditions, oxidant-stress derived) odd chain acyl-carnitines (e.g., undecanoyl-carnitine) was associated with polymorphisms in the coding region of the gene EPHX2, which was in turn also associated with variance in the levels of several linoleyl-derived oxylipins (9,10-EpOME, 12,13-EpOME), which are lipid mediators released by RBCs in response to hypoxia.^80^ It is worth noting that the levels of arachidonic acid were here found to associate with polymorphisms in fatty acid desaturase 1, confirming prior findings.^81^ This is relevant in that mature RBCs have been found to express functional fatty acid desaturases (FADS), and FADS activity – which is dependent on iron - was found to increase in response to storage-induced or pathological oxidant stress in vitro and in vivo.^82^ Prior work in mice found an association between the levels of oxylipins, iron metabolism (the ferrireductase STEAP3) and poor post-transfusion recovery of stored RBCs.^83^ Of note, another iron-dependent enzyme,^84^ spermine oxidase (SMOX) was found to be polymorphic in routine blood donors, which was here associated with varying levels of the product of its enzymatic activity, the polyamine spermine.

One of the main findings of the genomic arm of the REDS-III RBC-Omics Study was the identification of polymorphisms associated with the expression of a less active isoform of G6PD (African variant) that are associated with an increased susceptibility to end of storage hemolysis of RBCs following oxidant insults.^28^ Parallel metabolomics studies identified an impact for donor sex, ethnicity and age on the antioxidant systems (especially glutathione-dependent systems) of stored RBCs, with an emphasis for an impairment in the stored RBC capacity to activate the pentose phosphate pathway^16^ (G6PD is the rate-limiting enzyme of this pathway). These results were independently corroborated by the observation that failure to activate the PPP is a hallmark of the metabolic lesion to stored RBCs, in part attributable to the inability to inhibit glycolysis via the reversible oxidation of glyceraldehyde 3-phosphate dehydrogenase (GAPDH)^85^ and loss of GAPDH binding to the N-terminus cytosolic domain of band 3, ^86,87^ owing to fragmentation of the latter, as mediated by caspase activity or oxidant stress.^88^ Here we report that the levels of pyruvate in fresh RBCs and pyruvate/lactate ratios in stored RBCs are associated with the same G6PD polymorphisms. A causal role of this correlation is established in humanized murine models of G6PD deficiency, since the same metabolic change is seen in RBCs that differ only in their form of G6PD (African or Mediterranean^64^ variants vs. nondeficient human form). These observations could be partly explained by the compensatory over-activation of NADH-dependent methemoglobin reductase to cope with increased oxidant stress in G6PD deficient erythrocytes.^31^ Indeed, methemoglobin reductase would compete with lactate dehydrogenase for NADH, rendering the enzymatic step of lactic fermentation to regenerate NADH back to NAD+ no longer critical to preserve glycolytic fluxes (NAD+ is an essential cofactor for GAPDH activity upstream to pyruvate and ATP synthesis in glycolysis). The G6PD African variant in this donor population was also linked to variance in the levels of dopamine, confirming previous biomarker analyses from the metabolome of G6PD deficient vs sufficient blood donors^16^. This is interesting in that monoamine oxidase-dependent dopamine synthesis is an NADPH-dependent process, with implications relevant to exercise physiology (e.g., the sense of well-being/stimulatory effect after exercise^89^).

The present study has several limitations. First, mQTL analyses were determined based upon genomic characterization of a cohort of volunteer routine blood donors. As such, disease-related polymorphisms that would be identified in cohorts of non-healthy patients (i.e., from persons who are ineligible to donate blood) would be intrinsically not amenable to identification as a result of our study design. On the other hand, while sufficiently healthy to donate blood, the donor population enrolled in this study also includes phenotypes of potential clinical relevance to disease phenotypes (e.g., to cardiovascular and other disease risk factors, such as obesity,^27^ smoking,^23^ alcohol consumption^25^). As such, some of the genome-wide associations reported here (e.g., carnitine and SLC22A16) may be translationally relevant beyond transfusion medicine when interpreted in the context of markers relevant to specific diseases (e.g., carnitine metabolism and obesity^27^). Furthermore, only fresh (~10 day old – i.e. freshest samples available for this cohort) RBCs from volunteer donor volunteers were tested in this study. As such it is unclear whether the findings are relevant to transfusion medicine (e.g., genetic underpinning of metabolic heterogeneity in end of storage RBCs) or to physiological (e.g., hypoxia) or pathological conditions in which alterations to RBC metabolism are mechanistically relevant. Indeed, some metabolic markers of the RBC storage lesion only accumulate in end of storage units (e.g., hypoxanthine).^35^ Future studies will need to address this issue, by focusing on RBC samples stored for longer periods of time. Although the small (n=250) number of participants available still allowed for robust association discovery, a larger number of samples in more ancestrally diverse participants will increase the statistical power of future work. Such studies could pave the way for the use of other orthogonal omics approaches to metabolomics (e.g., proteomics) to maximize the value of the genetic and metabolic data already available for this well-curated cohort. Similar studies could be possible on other cohorts from patients with hematological conditions, such as sickle cell disease, where metabolite levels could not only be associated with, but also mechanistically contribute to the etiology of thromboinflammatory comorbidities of clinical relevance (e.g., sphingosine 1-phosphate and systemic hypoxemia,^90^ vasocclusive crisis, cardiopulmonary function, kidney dysfunction, pain crisis, etc.).

## Disclosure of Conflict of interest

Though unrelated to the contents of this manuscripts, the authors declare that AD is a founder of Omix Technologies Inc and Altis Biosciences LLC. AD is SAB members for Hemanext Inc and FORMA Therapeutics Inc. AD is a consultant for Rubius Therapeutics. JCZ is a consultant for Rubius Therapeutics and a founder of Svalinn Therapeutics. All other authors have no conflicts of interests to disclose.

## Acknowledgments

Research reported in this publication was funded by the National Institute of General and Medical Sciences (RM1GM131968 to ADA), and R01HL146442 (ADA), R01HL149714 (ADA), R01HL148151 (ADA), R01HL161004 (ADA), and R21HL150032 (ADA) from the National Heart, Lung, and Blood Institute. The content is solely the responsibility of the authors and does not necessarily represent the official views of the National Institutes of Health. The authors wish to acknowledge NHLBI Recipient Epidemiology and Donor Evaluation Study-III (REDS-III), which was supported by NHLBI contracts NHLBI NHLBI HHSN2682011-00001I, -00002I, -00003I, -00004I, -00005I, -00006I, -00007I, -00008I, and -00009I. The authors would like to express their deep gratitude Dr. Simone Glynn of NHLBI for her outstanding support throughout this study, the RBC-Omics research staff at all participating blood centers and testing labs for their exceptional performance and contribution to this project, and to all blood donors who agreed to participate in this study.

## The NHLBI Recipient Epidemiology Donor Evaluation Study-III (REDS-III), Red Blood Cell (RBC)-Omics Study, is the responsibility of the following persons

### Hubs

- Versiti Milwaukee, WI: A.E. Mast, J.L. Gottschall, W. Bialkowski, L. Anderson, J. Miller, A. Hall, Z. Udee, V. Johnson
- The Institute for Transfusion Medicine (ITXM), Pittsburgh, PA: D.J. Triulzi, J.E. Kiss, P.A. D’Andrea
- University of California, San Francisco, San Francisco, CA: E.L. Murphy, A.M. Guiltinan
- American Red Cross Blood Services, Farmington, CT: R.G. Cable, B.R. Spencer, S.T. Johnson
- Data coordinating center: RTI International, Rockville, MD: D.J. Brambilla, M.T. Sullivan, S.M. Endres-Dighe, G.P. Page, Y. Guo, N. Haywood, D. Ringer, B.C. Siege
- Central and testing laboratories: Vitalant Research Institute, San Francisco, CA: M.P. Busch, M.C. Lanteri, M. Stone, S. Keating
- Pittsburgh Heart, Lung, Blood, and Vascular Medicine Institute, Division of Pulmonary, Allergy and Critical Care Medicine, University of Pittsburgh, Pittsburgh, PA: T. Kanias, M. Gladwin
- Steering committee chairman: University of British Columbia, Victoria, BC, Canada: S.H. Kleinman
- National Heart, Lung, and Blood Institute, National Institutes of Health: S.A. Glynn, K.B. Malkin, A.M. Cristman

## Authors’ contributions

MPB led the REDS-III RBC-Omics project. AM and GPP performed mQTL analyses. AD performed metabolomics analyses. EAH and ROF co-ordinated human G6PD studies. KD and JCZ performed mouse studies. AM, AD, GP prepared figures and tables. AD wrote the first version of the manuscript and all authors contributed to its finalization.

## References

1. Nemkov T, Reisz JA, Xia Y, Zimring JC, D’Alessandro A. Red blood cells as an organ? How deep omics characterization of the most abundant cell in the human body highlights other systemic metabolic functions beyond oxygen transport. Expert Rev Proteomics. 2018;15(11):855–864.

2. D’Alessandro A, Dzieciatkowska M, Nemkov T, Hansen KC. Red blood cell proteomics update: is there more to discover? Blood Transfus. 2017;15(2):182–187.

3. Bryk AH, Wiśniewski JR. Quantitative Analysis of Human Red Blood Cell Proteome. J Proteome Res. 2017;16(8):2752–2761.

4. Wilson MC, Trakarnsanga K, Heesom KJ, et al. Comparison of the Proteome of Adult and Cord Erythroid Cells, and Changes in the Proteome Following Reticulocyte Maturation. Mol Cell Proteomics. 2016;15(6):1938–1946.

5. Kaestner L, Minetti G. The potential of erythrocytes as cellular aging models. Cell Death Differ. 2017;24(9):1475–1477.

6. D’Alessandro A, Giardina B, Gevi F, Timperio AM, Zolla L. Clinical metabolomics: the next stage of clinical biochemistry. Blood Transfus. 2012;10 Suppl 2(Suppl 2):s19–24.

7. Jacob M, Lopata AL, Dasouki M, Abdel Rahman AM. Metabolomics toward personalized medicine. Mass Spectrom Rev. 2019;38(3):221–238.

8. Zampieri M, Sekar K, Zamboni N, Sauer U. Frontiers of high-throughput metabolomics. Curr Opin Chem Biol. 2017;36:15–23.

9. D’Alessandro A, D’Amici GM, Vaglio S, Zolla L. Time-course investigation of SAGM-stored leukocyte-filtered red bood cell concentrates: from metabolism to proteomics. Haematologica. 2012;97(1):107–115.

10. Roussel C, Morel A, Dussiot M, et al. Rapid clearance of storage-induced microerythrocytes alters transfusion recovery. Blood. 2021;137(17):2285–2298.

11. Yoshida T, Prudent M, D’Alessandro A. Red blood cell storage lesion: causes and potential clinical consequences. Blood Transfus. 2019;17(1):27–52.

12. Roubinian NH, Plimier C, Woo JP, et al. Effect of donor, component, and recipient characteristics on hemoglobin increments following red blood cell transfusion. Blood. 2019;134(13):1003–1013.

13. Paglia G, D’Alessandro A, Rolfsson O, et al. Biomarkers defining the metabolic age of red blood cells during cold storage. Blood. 2016;128(13):e43–50.

14. Bordbar A, Johansson PI, Paglia G, et al. Identified metabolic signature for assessing red blood cell unit quality is associated with endothelial damage markers and clinical outcomes. Transfusion. 2016;56(4):852–862.

15. Van ’t Erve TJ, Wagner BA, Martin SM, et al. The heritability of hemolysis in stored human red blood cells. Transfusion. 2015;55(6):1178–1185.

16. D’Alessandro A, Fu X, Kanias T, et al. Donor sex, age and ethnicity impact stored red blood cell antioxidant metabolism through mechanisms in part explained by glucose 6-phosphate dehydrogenase levels and activity. Haematologica. 2021;106(5):1290–1302.

17. Francis RO, D’Alessandro A, Eisenberger A, et al. Donor glucose-6-phosphate dehydrogenase deficiency decreases blood quality for transfusion. J Clin Invest. 2020;130(5):2270–2285.

18. D’Alessandro A, Yoshida T, Nestheide S, et al. Hypoxic storage of red blood cells improves metabolism and post-transfusion recovery. Transfusion. 2020;60(4):786–798.

19. Donovan K, Meli A, Cendali F, et al. Stored blood has compromised oxygen unloading kinetics that can be normalized with rejuvenation and predicted from corpuscular side-scatter. Haematologica. 2021.

20. D’Alessandro A, Zimring JC, Busch M. Chronological storage age and metabolic age of stored red blood cells: are they the same? Transfusion. 2019;59(5):1620–1623.

21. D’Alessandro A, Fu X, Reisz JA, et al. Stored RBC metabolism as a function of caffeine levels. Transfusion. 2020;60(6):1197–1211.

22. Nemkov T, Stefanoni D, Bordbar A, et al. Blood donor exposome and impact of common drugs on red blood cell metabolism. JCI Insight. 2020.

23. Stefanoni D, Fu X, Reisz JA, et al. Nicotine exposure increases markers of oxidant stress in stored red blood cells from healthy donor volunteers. Transfusion. 2020;60(6):1160–1174.

24. DeSimone RA, Hayden JA, Mazur CA, et al. Red blood cells donated by smokers: A pilot investigation of recipient transfusion outcomes. Transfusion. 2019;59(8):2537–2543.

25. D’Alessandro A, Fu X, Reisz JA, et al. Ethyl glucuronide, a marker of alcohol consumption, correlates with metabolic markers of oxidant stress but not with hemolysis in stored red blood cells from healthy blood donors. Transfusion. 2020;60(6):1183–1196.

26. Kanias T, Lanteri MC, Page GP, et al. Ethnicity, sex, and age are determinants of red blood cell storage and stress hemolysis: results of the REDS-III RBC-Omics study. Blood Adv. 2017;1(15):1132–1141.

27. Hazegh K, Fang F, Bravo MD, et al. Blood donor obesity is associated with changes in red blood cell metabolism and susceptibility to hemolysis in cold storage and in response to osmotic and oxidative stress. Transfusion. 2021;61(2):435–448.

28. Page GP, Kanias T, Guo YJ, et al. Multiple-ancestry genome-wide association study identifies 27 loci associated with measures of hemolysis following blood storage. J Clin Invest. 2021;131(13).

29. D’Alessandro A, Culp-Hill R, Reisz JA, et al. Heterogeneity of blood processing and storage additives in different centers impacts stored red blood cell metabolism as much as storage time: lessons from REDS-III-Omics. Transfusion. 2019;59(1):89–100.

30. Guo Y, Busch MP, Seielstad M, et al. Development and evaluation of a transfusion medicine genome wide genotyping array. Transfusion. 2019;59(1):101–111.

31. Tzounakas VL, Kriebardis AG, Georgatzakou HT, et al. Glucose 6-phosphate dehydrogenase deficient subjects may be better “storers” than donors of red blood cells. Free Radic Biol Med. 2016;96:152–165.

32. Roubinian NH, Reese SE, Qiao H, et al. Donor genetic and nongenetic factors affecting red blood cell transfusion effectiveness. JCI Insight. 2022;7(1).

33. D’Alessandro A, Hansen KC, Eisenmesser EZ, Zimring JC. Protect, repair, destroy or sacrifice: a role of oxidative stress biology in inter-donor variability of blood storage? Blood Transfus. 2019;17(4):281–288.

34. Burka ER, Weaver Z, 3rd, Marks PA. Clinical spectrum of hemolytic anemia associated with glucose-6-phosphate dehydrogenase deficiency. Ann Intern Med. 1966;64(4):817–825.

35. Nemkov T, Sun K, Reisz JA, et al. Hypoxia modulates the purine salvage pathway and decreases red blood cell and supernatant levels of hypoxanthine during refrigerated storage. Haematologica. 2018;103(2):361–372.

36. Ingrosso D, Cimmino A, D’Angelo S, Alfinito F, Zappia V, Galletti P. Protein methylation as a marker of aspartate damage in glucose-6-phosphate dehydrogenase-deficient erythrocytes: role of oxidative stress. Eur J Biochem. 2002;269(8):2032–2039.

37. Sabino EC, Buss LF, Carvalho MPS, et al. Resurgence of COVID-19 in Manaus, Brazil, despite high seroprevalence. The Lancet. 2021;397(10273):452–455.

38. Kraus WE, Muoio DM, Stevens R, et al. Metabolomic Quantitative Trait Loci (mQTL) Mapping Implicates the Ubiquitin Proteasome System in Cardiovascular Disease Pathogenesis. PLOS Genetics. 2015;11(11):e1005553.

39. Kurbatova N, Garg M, Whiley L, et al. Urinary metabolic phenotyping for Alzheimer’s disease. Scientific Reports. 2020;10(1):21745.

40. Panyard DJ, Kim KM, Darst BF, et al. Cerebrospinal fluid metabolomics identifies 19 brain-related phenotype associations. Communications biology. 2021;4(1):63–63.

41. Yousri NA, Fakhro KA, Robay A, et al. Whole-exome sequencing identifies common and rare variant metabolic QTLs in a Middle Eastern population. Nat Commun. 2018;9(1):333.

42. Johnson RK, Brunetti T, Quinn K, et al. Discovering metabolite quantitative trait loci in asthma using an isolated population. J Allergy Clin Immunol. 2021.

43. Nemkov T, Skinner SC, Nader E, et al. Acute Cycling Exercise Induces Changes in Red Blood Cell Deformability and Membrane Lipid Remodeling. Int J Mol Sci. 2021;22(2).

44. Dong S, Wang Q, Kao YR, et al. Chaperone-mediated autophagy sustains haematopoietic stem-cell function. Nature. 2021;591(7848):117–123.

45. D’Alessandro A, Nemkov T, Sun K, et al. AltitudeOmics: Red Blood Cell Metabolic Adaptation to High Altitude Hypoxia. J Proteome Res. 2016;15(10):3883–3895.

46. Reisz J, Slaughter A, D’Alessandro A, al. e. Red blood cells in hemorrhagic shock: a critical role for glutaminolysis in fueling alanine transamination in rats. blood advances. 2017;1(17):1296–1305.

47. Thomas T, Stefanoni D, Dzieciatkowska M, et al. Evidence of Structural Protein Damage and Membrane Lipid Remodeling in Red Blood Cells from COVID-19 Patients. J Proteome Res. 2020;19(11):4455–4469.

48. Renoux C, Fort R, Nader E, et al. Impact of COVID-19 on red blood cell rheology. Br J Haematol. 2021;192(4):e108–e111.

49. Pernow J, Mahdi A, Yang J, Zhou Z. Red blood cell dysfunction: a new player in cardiovascular disease. Cardiovascular Research. 2019;115(11):1596–1605.

50. Bissinger R, Nemkov T, D’Alessandro A, et al. Proteinuric chronic kidney disease is associated with altered red blood cell lifespan, deformability and metabolism. Kidney Int. 2021;100(6):1227–1239.

51. Xu P, Chen C, Zhang Y, et al. Erythrocyte transglutaminase-2 combats hypoxia and chronic kidney disease by promoting oxygen delivery and carnitine homeostasis. Cell Metabolism. 2022;34(2):299–316.e296.

52. Nemkov T, Reisz JA, Gehrke S, Hansen KC, D’Alessandro A. High-Throughput Metabolomics: Isocratic and Gradient Mass Spectrometry-Based Methods. Methods Mol Biol. 2019;1978:13–26.

53. Reisz JA, Zheng C, D’Alessandro A, Nemkov T. Untargeted and Semi-targeted Lipid Analysis of Biological Samples Using Mass Spectrometry-Based Metabolomics. Methods Mol Biol. 2019;1978:121–135.

54. Nemkov T, Hansen KC, D’Alessandro A. A three-minute method for high-throughput quantitative metabolomics and quantitative tracing experiments of central carbon and nitrogen pathways. Rapid Commun Mass Spectrom. 2017;31(8):663–673.

55. D’Alessandro A, Culp-Hill R, Reisz JA, et al. Heterogeneity of blood processing and storage additives in different centers impacts stored red blood cell metabolism as much as storage time: lessons from REDS-III-Omics. Transfusion. 2019;59(1):89–100.

56. Wei R, Wang J, Su M, et al. Missing Value Imputation Approach for Mass Spectrometry-based Metabolomics Data. Scientific Reports. 2018;8(1):663.

57. Ongen H, Buil A, Brown AA, Dermitzakis ET, Delaneau O. Fast and efficient QTL mapper for thousands of molecular phenotypes. Bioinformatics. 2015;32(10):1479–1485.

58. Delaneau O, Coulonges C, Zagury J-F. Shape-IT: new rapid and accurate algorithm for haplotype inference. BMC Bioinformatics. 2008;9(1):540.

59. Howie B, Marchini J, Stephens M. Genotype Imputation with Thousands of Genomes. G3 Genes|Genomes|Genetics. 2011;1(6):457–470.

60. Zheng X, Levine D, Shen J, Gogarten SM, Laurie C, Weir BS. A high-performance computing toolset for relatedness and principal component analysis of SNP data. Bioinformatics. 2012;28(24):3326–3328.

61. Aulchenko YS, Struchalin MV, van Duijn CM. ProbABEL package for genome-wide association analysis of imputed data. BMC Bioinformatics. 2010;11(1):134.

62. Perry JA, Gaynor BJ, Mitchell BD, O’Connell JR. An Omics Analysis Search and Information System (OASIS) for Enabling Biological Discovery in the Old Order Amish. bioRxiv. 2021:2021.2005.2002.442370.

63. Machiela MJ, Chanock SJ. LDlink: a web-based application for exploring population-specific haplotype structure and linking correlated alleles of possible functional variants. Bioinformatics. 2015;31(21):3555–3557.

64. D’Alessandro A, Howie HL, Hay AM, et al. Hematologic and systemic metabolic alterations due to Mediterranean class II G6PD deficiency in mice. JCI Insight. 2021;6(14).

65. Rueedi R, Ledda M, Nicholls A, et al. Genome-Wide Association Study of Metabolic Traits Reveals Novel Gene-Metabolite-Disease Links. PLoS genetics. 2014;10:e1004132.

66. Nicholson G, Rantalainen M, Li J, et al. A genome-wide metabolic QTL analysis in Europeans implicates two loci shaped by recent positive selection. PLoSGenet. 2011;7:e1002270.

67. Gillenwater LA, Pratte KA, Hobbs BD, et al. Plasma Metabolomic Signatures of Chronic Obstructive Pulmonary Disease and the Impact of Genetic Variants on Phenotype-Driven Modules. Network and systems medicine. 2020;3(1):159–181.

68. Olarini A, Ernst M, Gürdeniz G, et al. Vertical Transfer of Metabolites Detectable from Newborn’s Dried Blood Spot Samples Using UPLC-MS: A Chemometric Study. Metabolites. 2022;12(2).

69. Mercurio S, Petrillo S, Chiabrando D, et al. The heme exporter Flvcr1 regulates expansion and differentiation of committed erythroid progenitors by controlling intracellular heme accumulation. Haematologica. 2015;100(6):720–729.

70. Davis LJ, Brown A. THE ERYTHROPOIETIC ACTIVITY OF CHOLINE CHLORIDE IN MEGALOBLASTIC ANEMIAS. Blood. 1947;2(5):407–425.

71. D’Alessandro A, Hay A, Dzieciatkowska M, et al. Protein-L-isoaspartate O-methyltransferase is required for *in vivo* control of oxidative damage in red blood cells. Haematologica. 2021;106(10):2726–2739.

72. Reisz JA, Nemkov T, Dzieciatkowska M, et al. Methylation of protein aspartates and deamidated asparagines as a function of blood bank storage and oxidative stress in human red blood cells. Transfusion. 2018;58(12):2978–2991.

73. Huang N-J, Lin Y-C, Lin C-Y, et al. Enhanced phosphocholine metabolism is essential for terminal erythropoiesis. Blood. 2018;131(26):2955–2966.

74. Wang B, Tontonoz P. Phospholipid Remodeling in Physiology and Disease. Annual review of physiology. 2019;81:165–188.

75. Rong X, Wang B, Dunham MM, et al. Lpcat3-dependent production of arachidonoyl phospholipids is a key determinant of triglyceride secretion. eLife. 2015;4:e06557.

76. Khiati S, Dalla Rosa I, Sourbier C, et al. Mitochondrial topoisomerase I (top1mt) is a novel limiting factor of doxorubicin cardiotoxicity. Clinical cancer research : an official journal of the American Association for Cancer Research. 2014;20(18):4873–4881.

77. Bissinger R, Nemkov T, D’Alessandro A, et al. Proteinuric chronic kidney disease is associated with altered red blood cell lifespan, deformability and metabolism. Kidney International. 2021;100(6):1227–1239.

78. Wu H, Bogdanov M, Zhang Y, et al. Hypoxia-mediated impaired erythrocyte Lands’ Cycle is pathogenic for sickle cell disease. Sci Rep. 2016;6:29637.

79. Arduini A, Holme S, Sweeney JD, Dottori S, Sciarroni AF, Calvani M. Addition of L-carnitine to additive solution-suspended red cells stored at 4 degrees C reduces in vitro hemolysis and improves in vivo viability. Transfusion. 1997;37(2):166–174.

80. Jiang H, Anderson GD, McGiff JC. Red blood cells (RBCs), epoxyeicosatrienoic acids (EETs) and adenosine triphosphate (ATP). Pharmacol Rep. 2010;62(3):468–474.

81. Tintle NL, Pottala JV, Lacey S, et al. A genome-wide association study of saturated, mono-and polyunsaturated red blood cell fatty acids in the Framingham Heart Offspring Study. Prostaglandins, leukotrienes, and essential fatty acids. 2015;94:65–72.

82. Thomas T, Cendali F, Fu X, et al. Fatty acid desaturase activity in mature red blood cells and implications for blood storage quality. Transfusion. 2021.

83. Howie HL, Hay AM, de Wolski K, et al. Differences in Steap3 expression are a mechanism of genetic variation of RBC storage and oxidative damage in mice. Blood Adv. 2019;3(15):2272–2285.

84. Kalani Roy M, La Carpia F, Cendali F, et al. Irradiation Causes Alterations of Polyamine, Purine, and Sulfur Metabolism in Red Blood Cells and Multiple Organs. J Proteome Res. 2022;21(2):519–534.

85. Reisz JA, Wither MJ, Dzieciatkowska M, et al. Oxidative modifications of glyceraldehyde 3-phosphate dehydrogenase regulate metabolic reprogramming of stored red blood cells. Blood. 2016;128(12):e32–42.

86. Issaian A, Hay A, Dzieciatkowska M, et al. The interactome of the N-terminus of band 3 regulates red blood cell metabolism and storage quality. Haematologica. 2021.

87. Rogers SC, Ge X, Brummet M, et al. Quantifying dynamic range in red blood cell energetics: Evidence of progressive energy failure during storage. Transfusion. 2021;61(5):1586–1599.

88. Rinalducci S, Ferru E, Blasi B, Turrini F, Zolla L. Oxidative stress and caspase-mediated fragmentation of cytoplasmic domain of erythrocyte band 3 during blood storage. Blood Transfus. 2012; 10 Suppl 2(Suppl 2):s55–62.

89. San-Millán I, Stefanoni D, Martinez JL, Hansen KC, D’Alessandro A, Nemkov T. Metabolomics of Endurance Capacity in World Tour Professional Cyclists. Front Physiol. 2020;11:578.

90. Sun K, D’Alessandro A, Ahmed MH, et al. Structural and Functional Insight of Sphingosine 1-Phosphate-Mediated Pathogenic Metabolic Reprogramming in Sickle Cell Disease. Scientific Reports. 2017;7(1):15281.

