## Supplementary Figures for "Genome-wide metabolite quantitative trait loci analysis (mQTL) in red blood cells from volunteer blood donors"

**Supplementary Table 1 – Please refer to the file *Supplementary Table 1.xlsx***

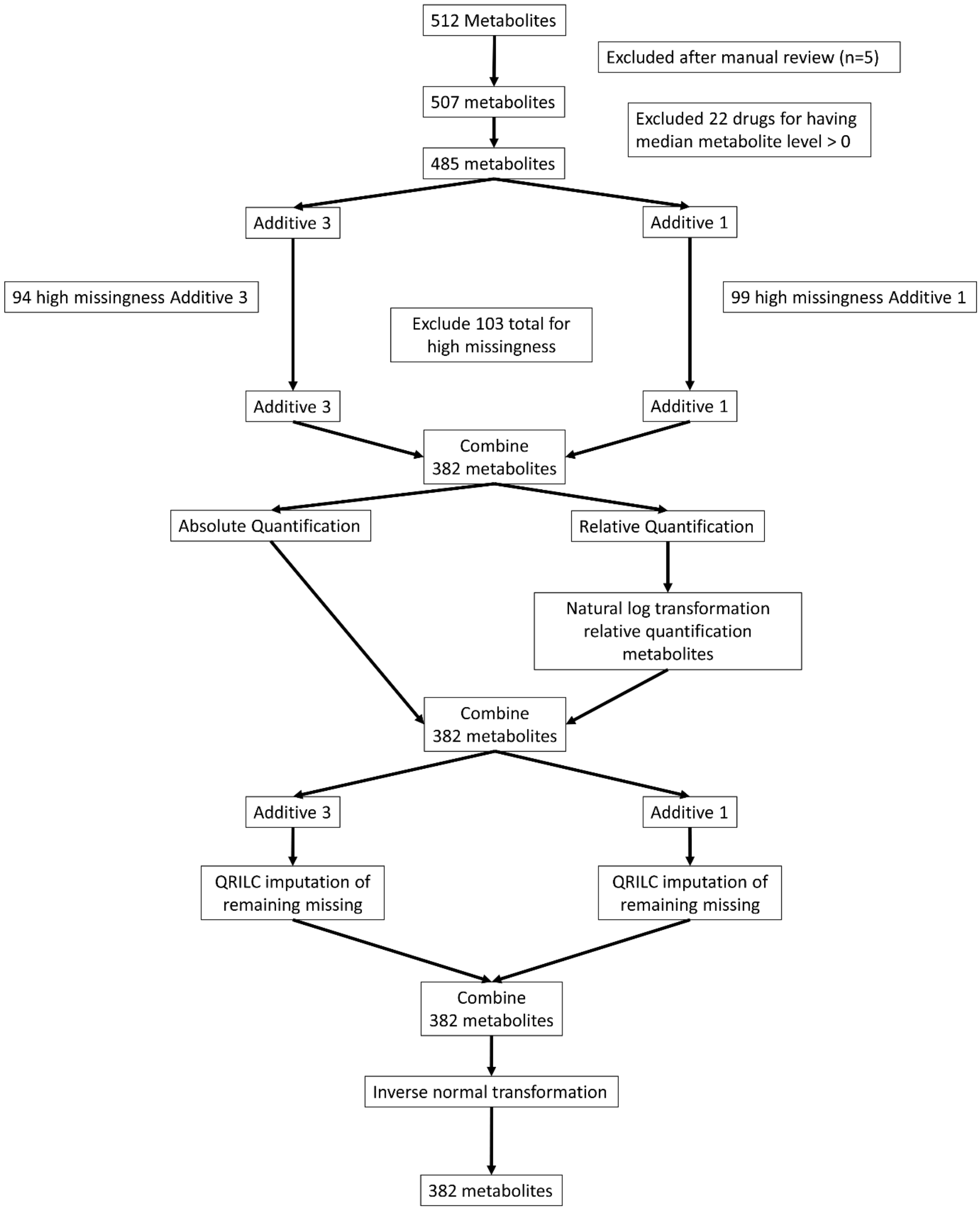


**Supplementary Figure 1 – Flow-chart of metabolomics post-processing.**

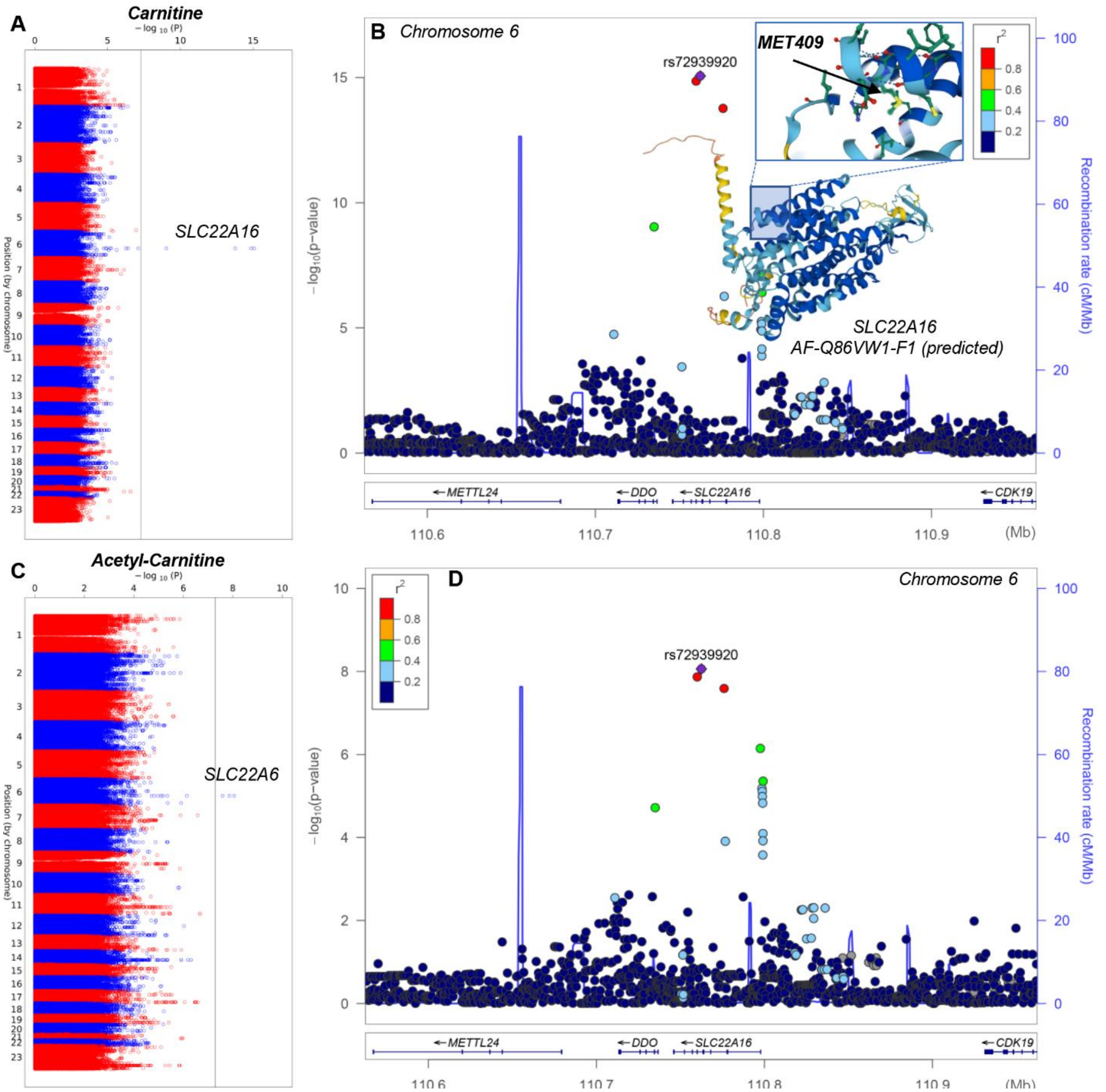


**Supplementary Figure 2 – Polymorphisms in the carnitine transporter SLC22A16 are associated with variability in the levels of RBC free and acetyl-carnitine.** Manhattan plots and LocusZooms are shown in panels **A-B** and **C-D**, respectively.

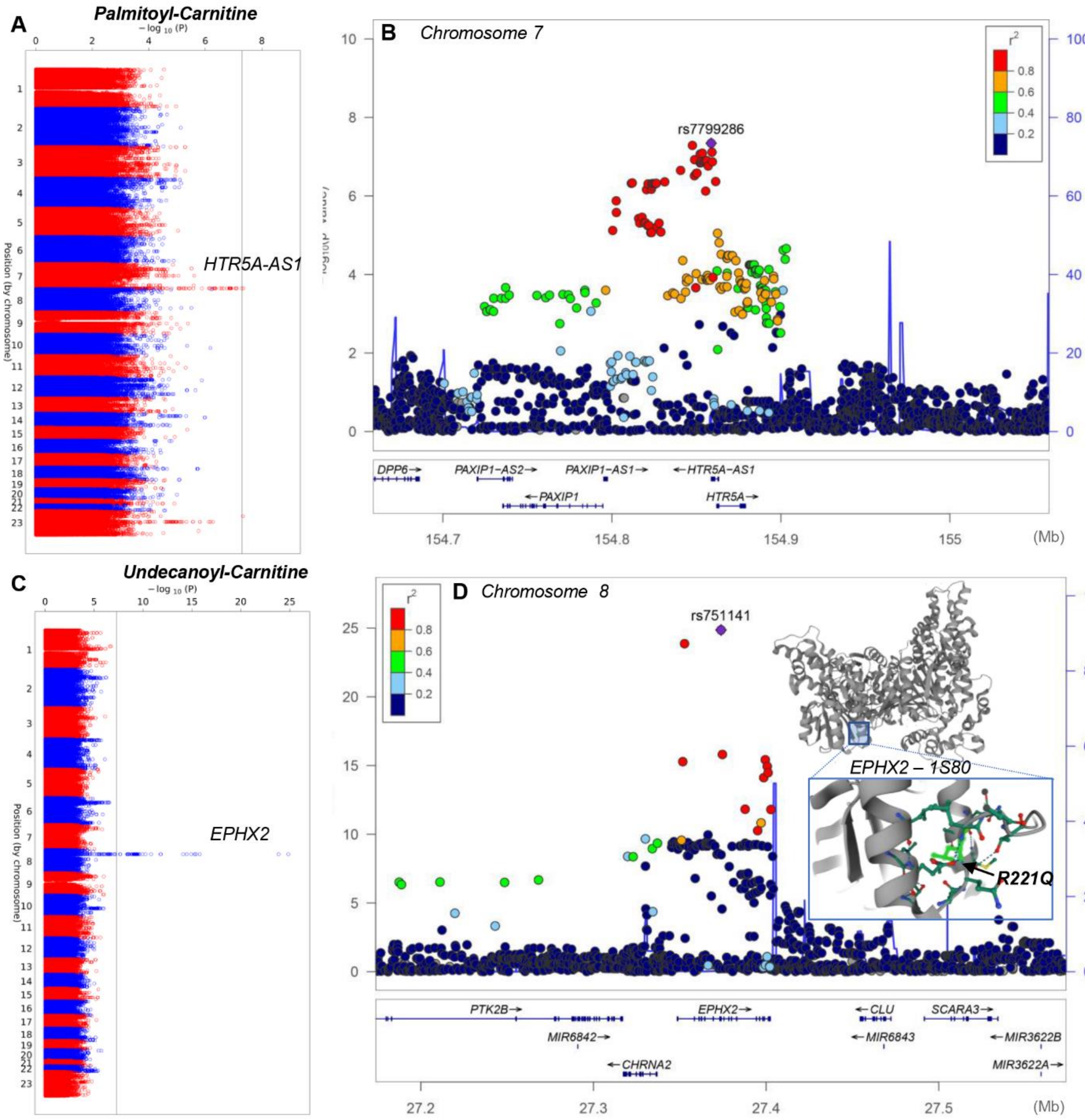


**Supplementary Figure 3 – Polymorphisms in HTR5A-AS1 and EPHX2 are associated with variability in the levels of RBC palmitoyl- and undecanoyl-carnitine- an odd-chain fatty acyl-carnitine derived from lipid peroxidation reactions.** Manhattan plots and LocusZooms are shown in panels **A-B** and **C-D**, respectively.

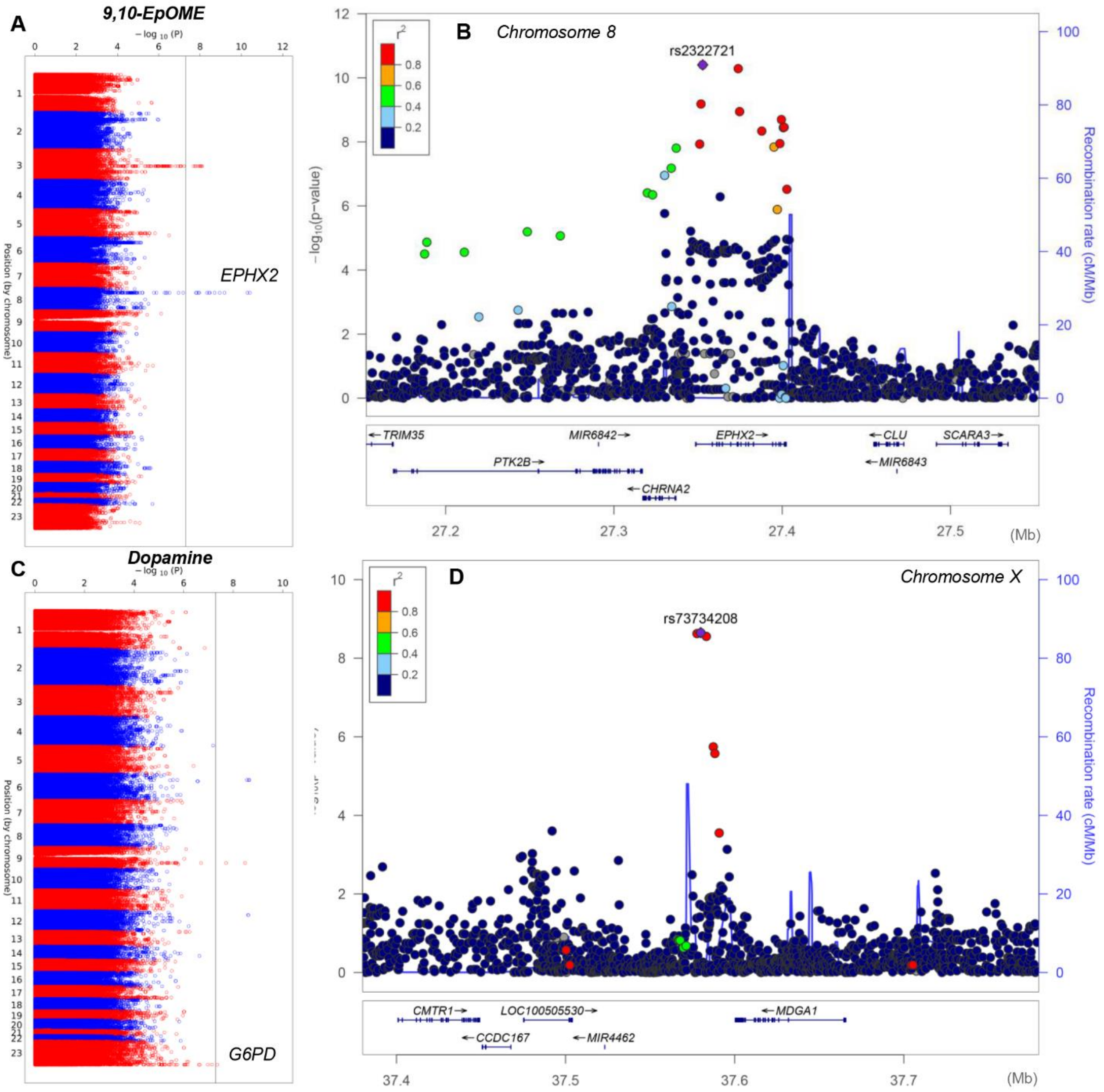


**Supplementary Figure 4 – Polymorphisms in the EPHX2 and G6PD are associated with variability in the levels of RBC 9,10,EpOME (oxylipin) and dopamine.** Manhattan plots and LocusZooms are shown in panels **A-B** and **C-D**, respectively.

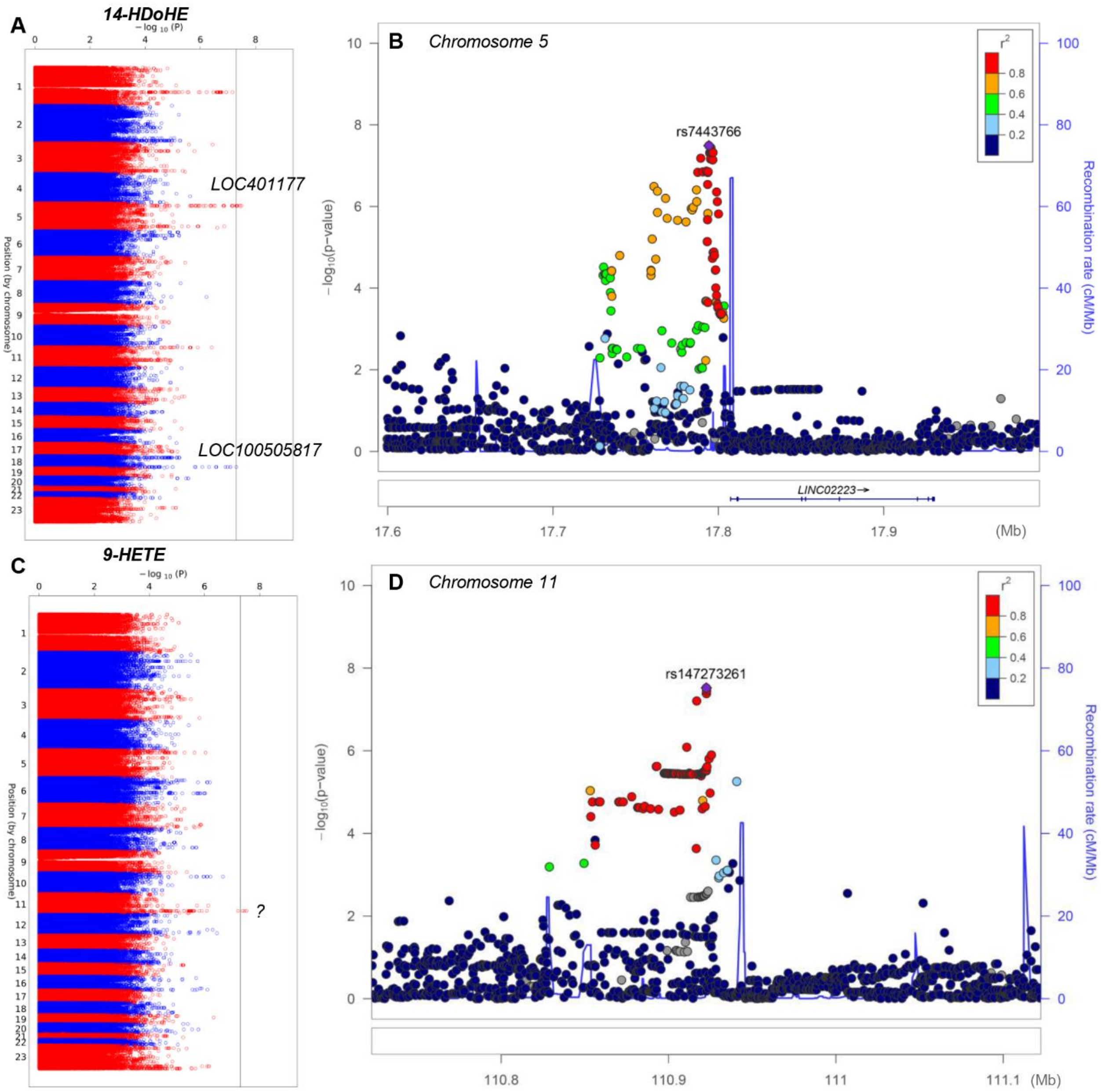


**Supplementary Figure 5 – Polymorphisms in LOC401177 and rs1472723261 SNP are associated with variability in the levels of RBC oxylipins 14-HDoHE and 9-HETE.** Manhattan plots and LocusZooms are shown in panels **A-B** and **C-D**, respectively.

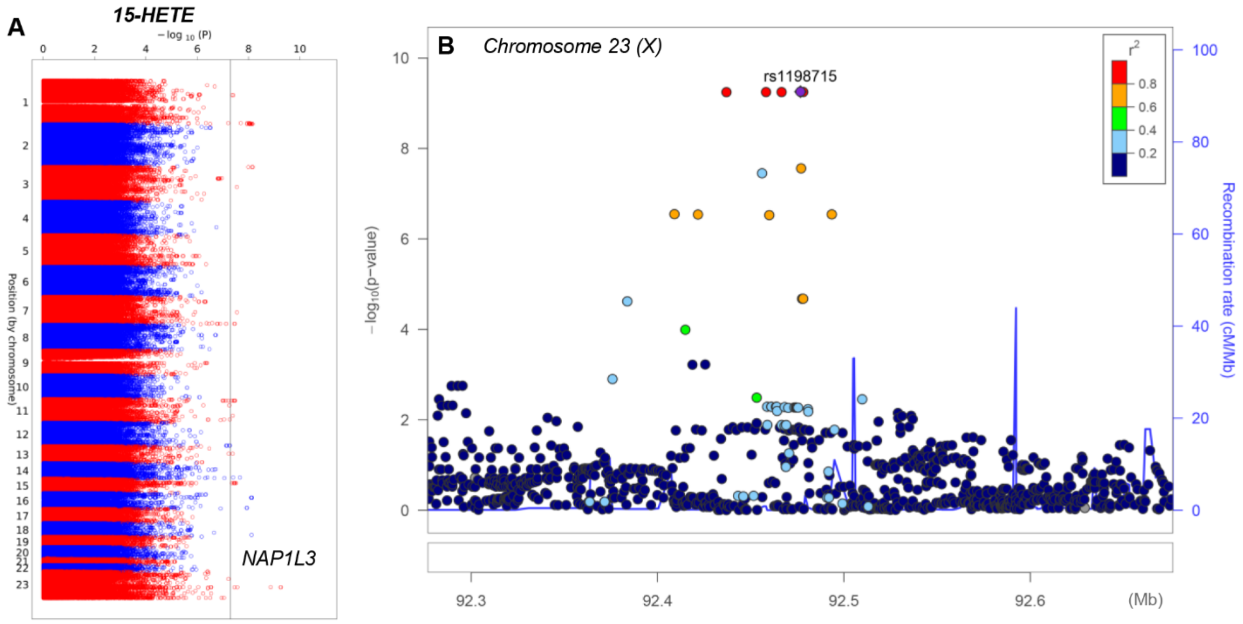


**Supplementary Figure 6 – Polymorphisms in the coding region for NAP1L3 are associated with variability in the levels of RBC 15-HETE.** Manhattan plots and LocusZooms are shown in panels **A-B** and **C-D**, respectively.

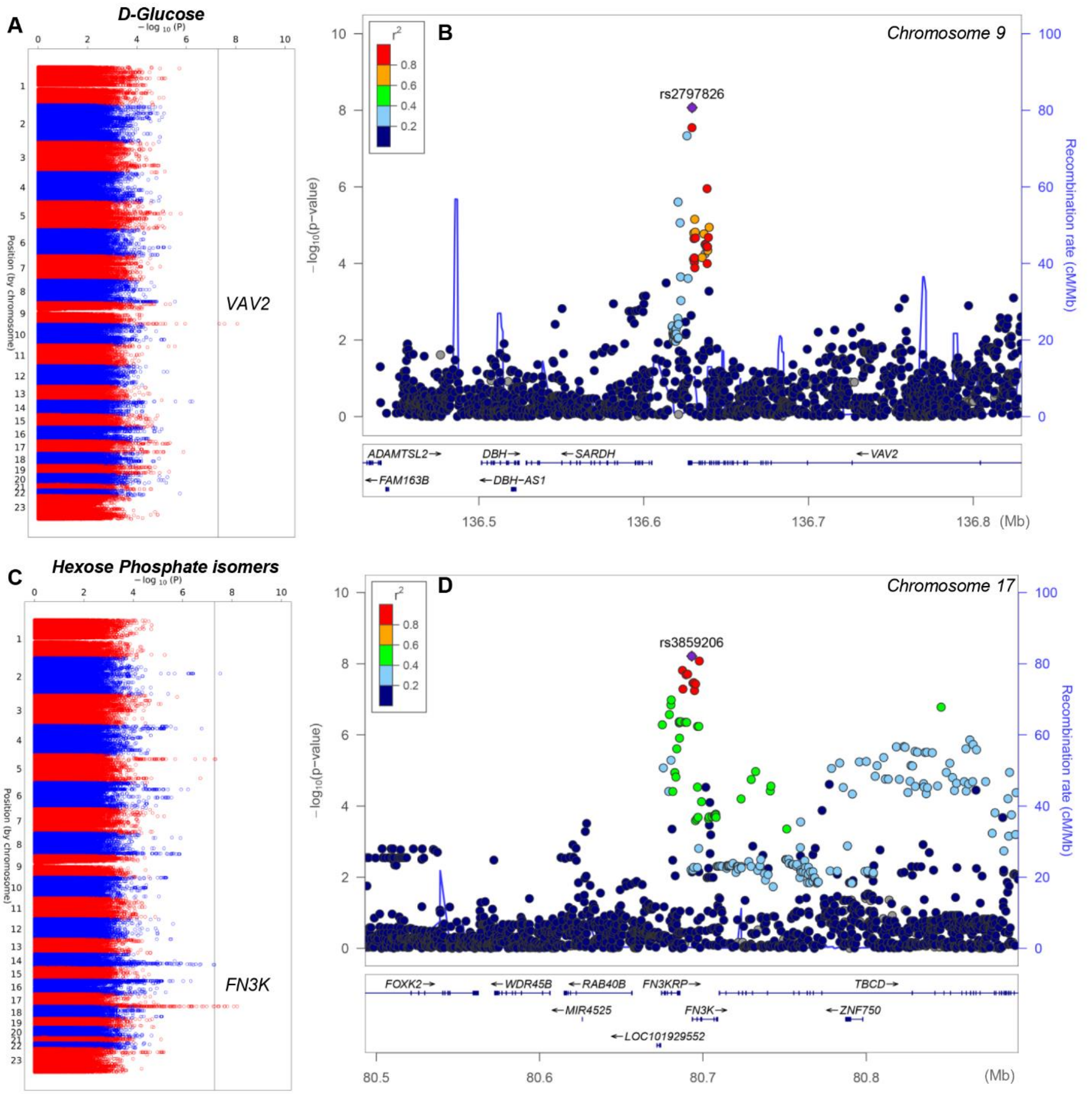


**Supplementary Figure 7 – Polymorphisms in VAV2 and FN3K are associated with variability in the levels of RBC glycolytic metabolites glucose and hexose phosphate isomers.** Manhattan plots and LocusZooms are shown in panels **A-B** and **C-D**, respectively.

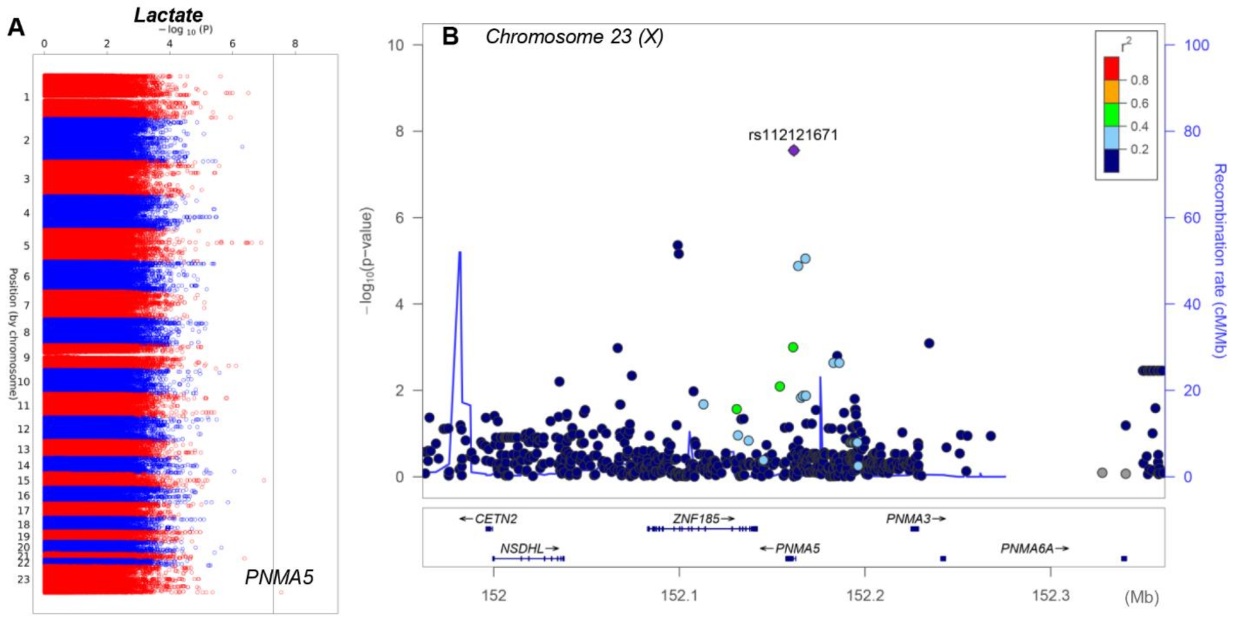


**Supplementary Figure 8 – Polymorphisms in PNMA5 are associated with variability in the levels of RBC lactate.** Manhattan plots and LocusZooms are shown in panels **A-B**, respectively.

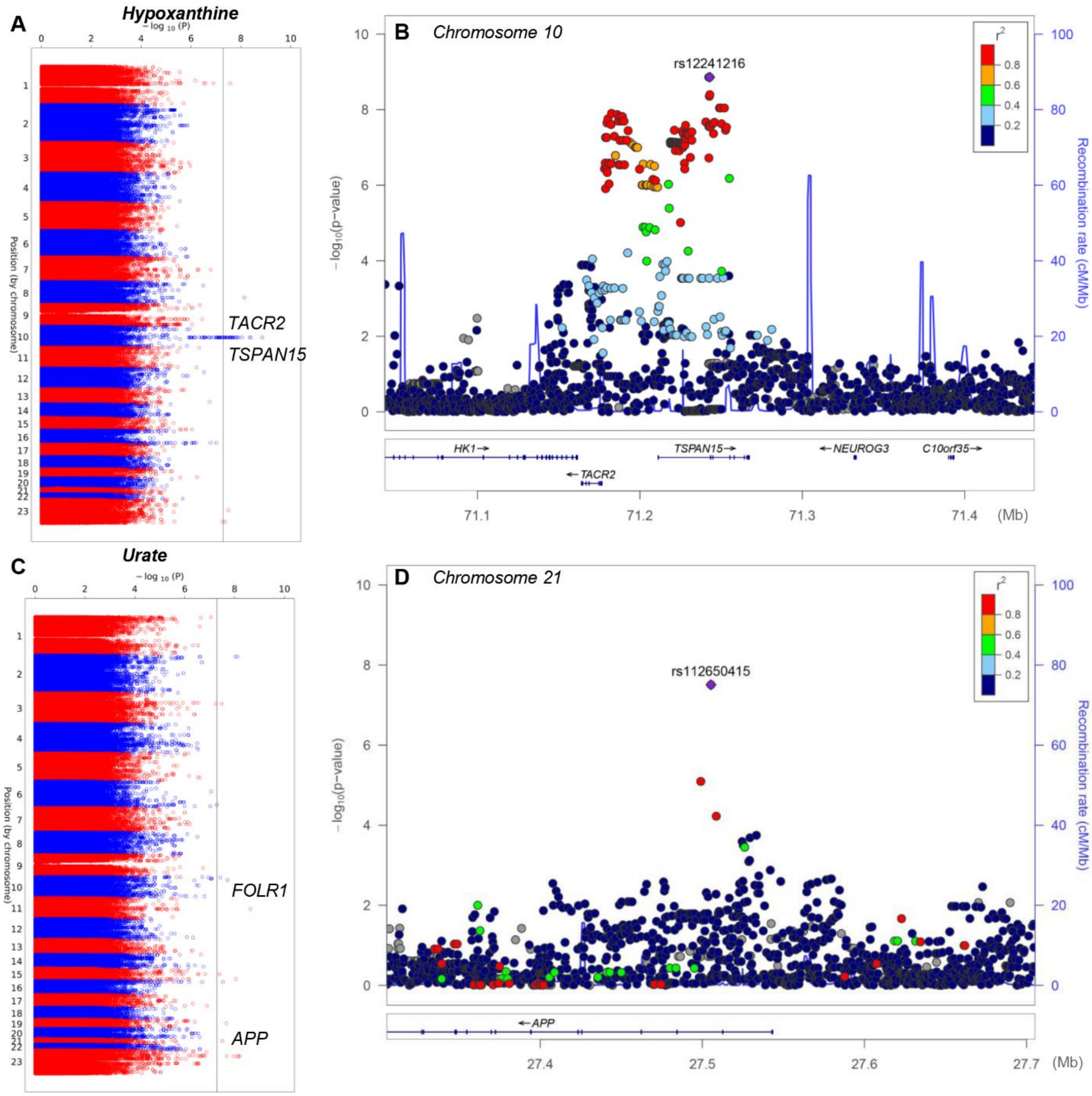


**Supplementary Figure 9 – Polymorphisms in TACR2 or TSPAN15 and APP or FOLR1 are associated with variability in the levels of RBC purine deamination products hypoxanthine and urate.** Manhattan plots and LocusZooms are shown in panels **A-B** and **C-D**, respectively.

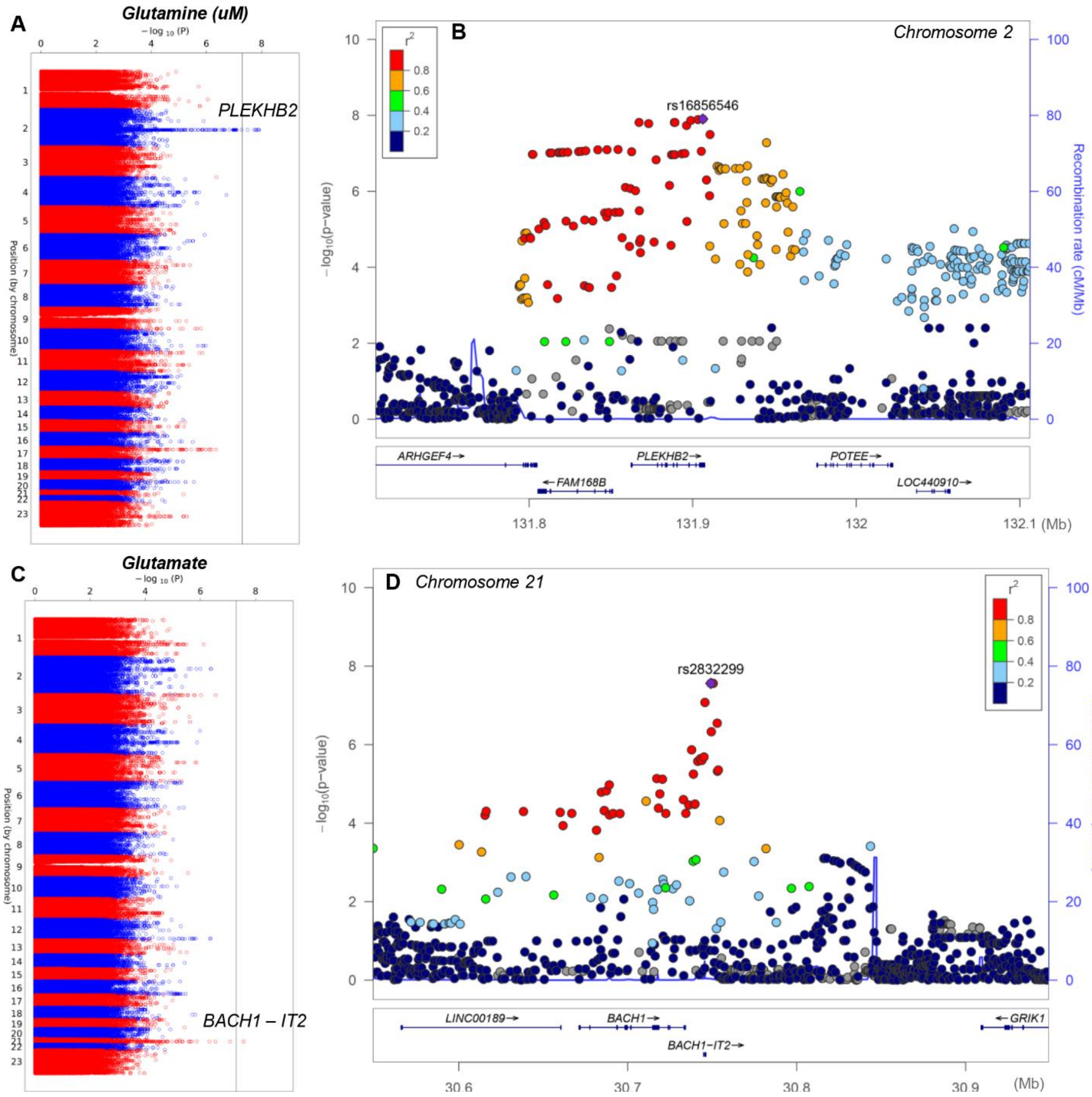


**Supplementary Figure 10 – Polymorphisms in PLEKHB2 and BACH1-IT2 are associated with variability in the levels of RBC amino acids glutamine and glutamate.** Manhattan plots and LocusZooms are shown in panels **A-B** and **C-D**, respectively.

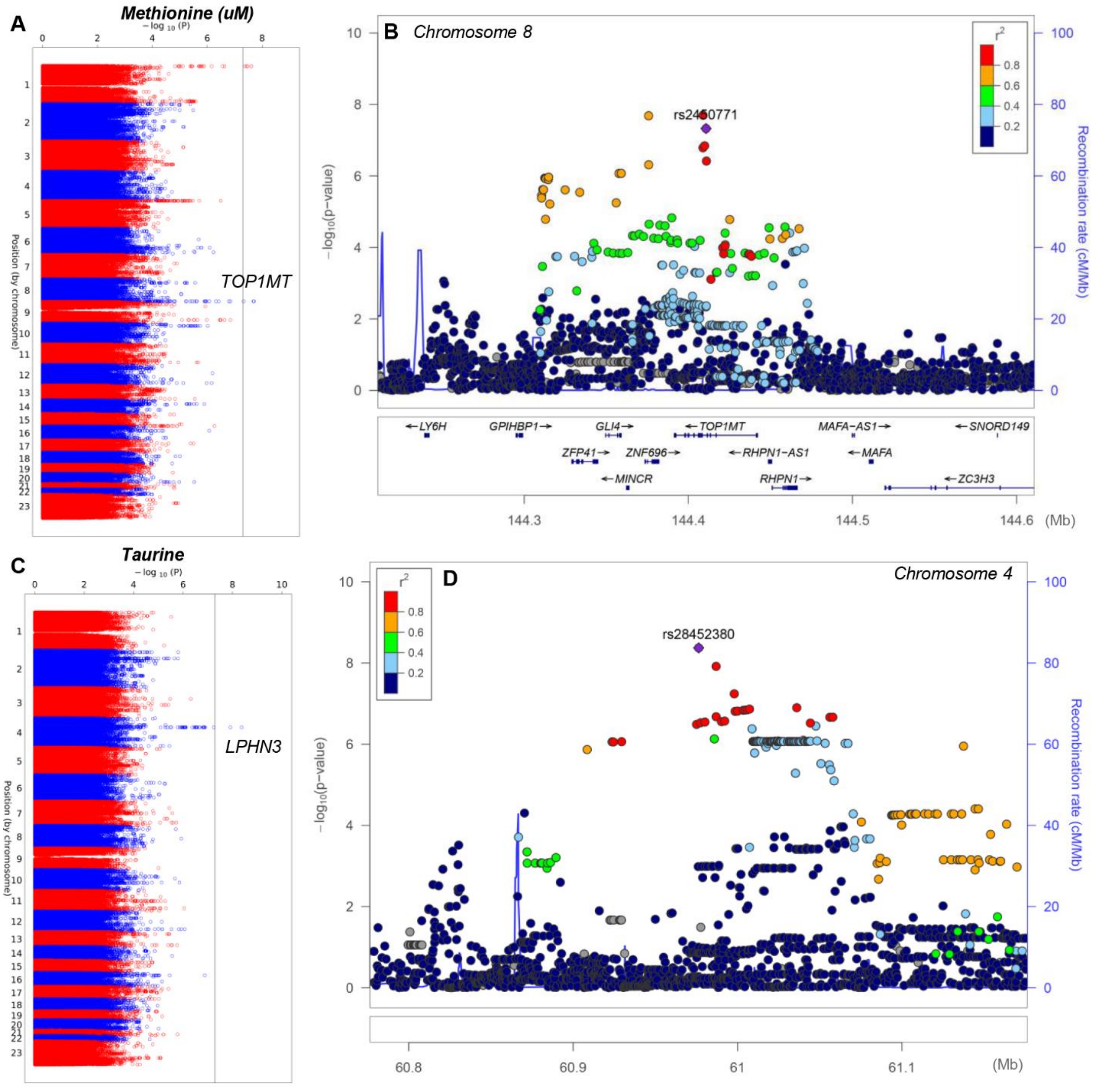


**Supplementary Figure 11 – Polymorphisms in TOP1MT and LPHN3 are associated with variability in the levels of RBC amino acids methionine and taurine.** Manhattan plots and LocusZooms are shown in panels **A-B** and **C-D**, respectively.

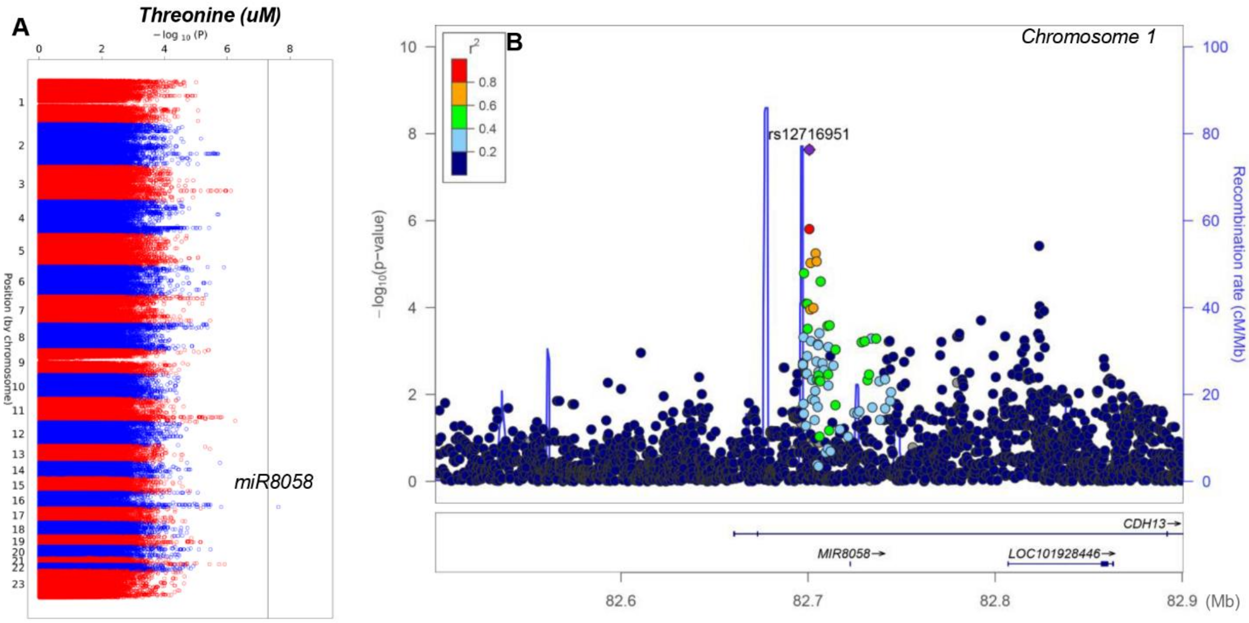


**Supplementary Figure 12 – Polymorphisms in miR8058 are associated with variability in the levels of RBC threonine.** Manhattan plots and LocusZooms are shown in panels **A-B**, respectively.

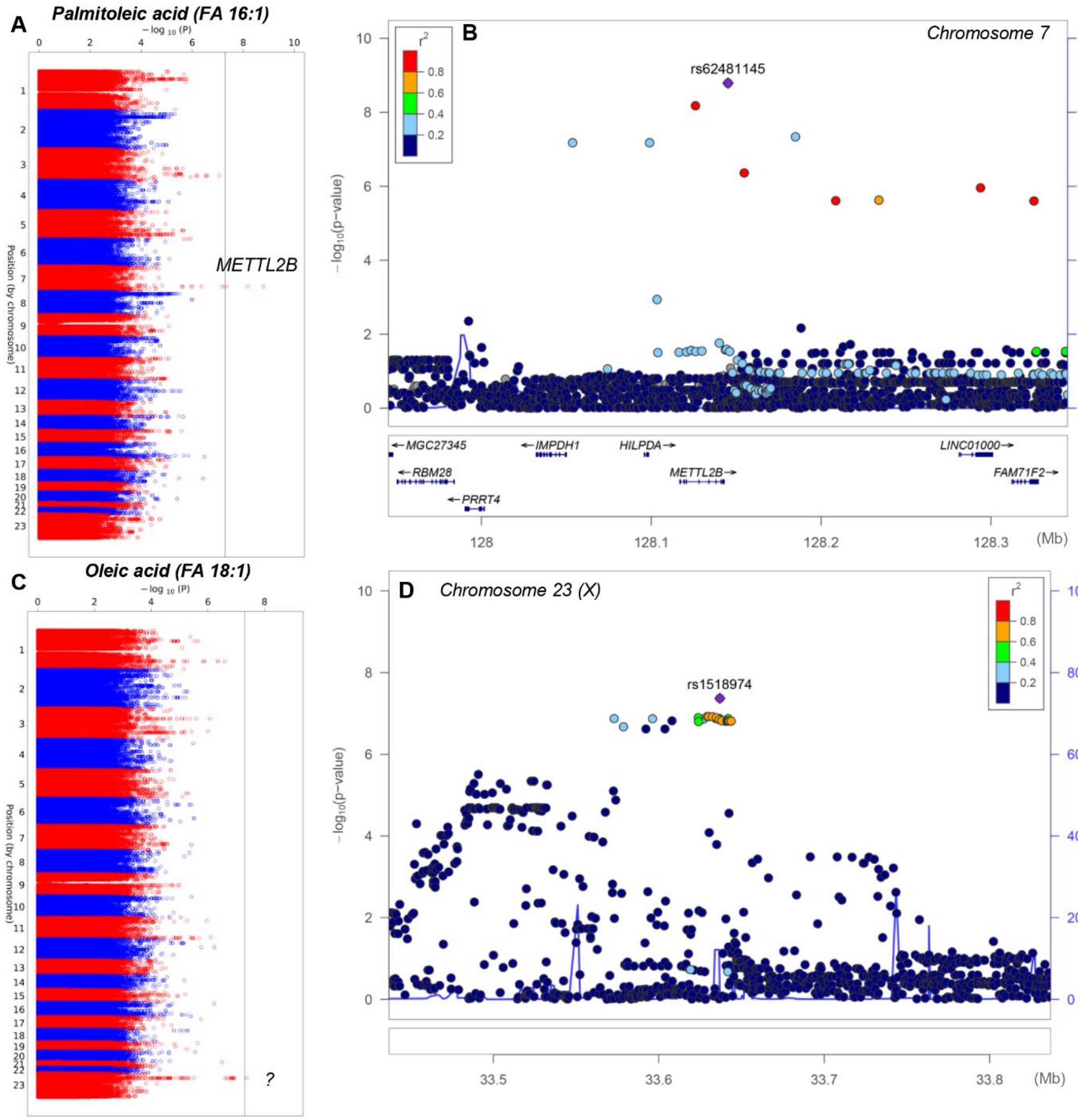


**Supplementary Figure 13 – Polymorphisms in METTL2b and rs1518974 SNP are associated with variability in the levels of RBC monounsaturated fatty acids palimitoleic and oleic acid.** Manhattan plots and LocusZooms are shown in panels **A-B** and **C-D**, respectively.

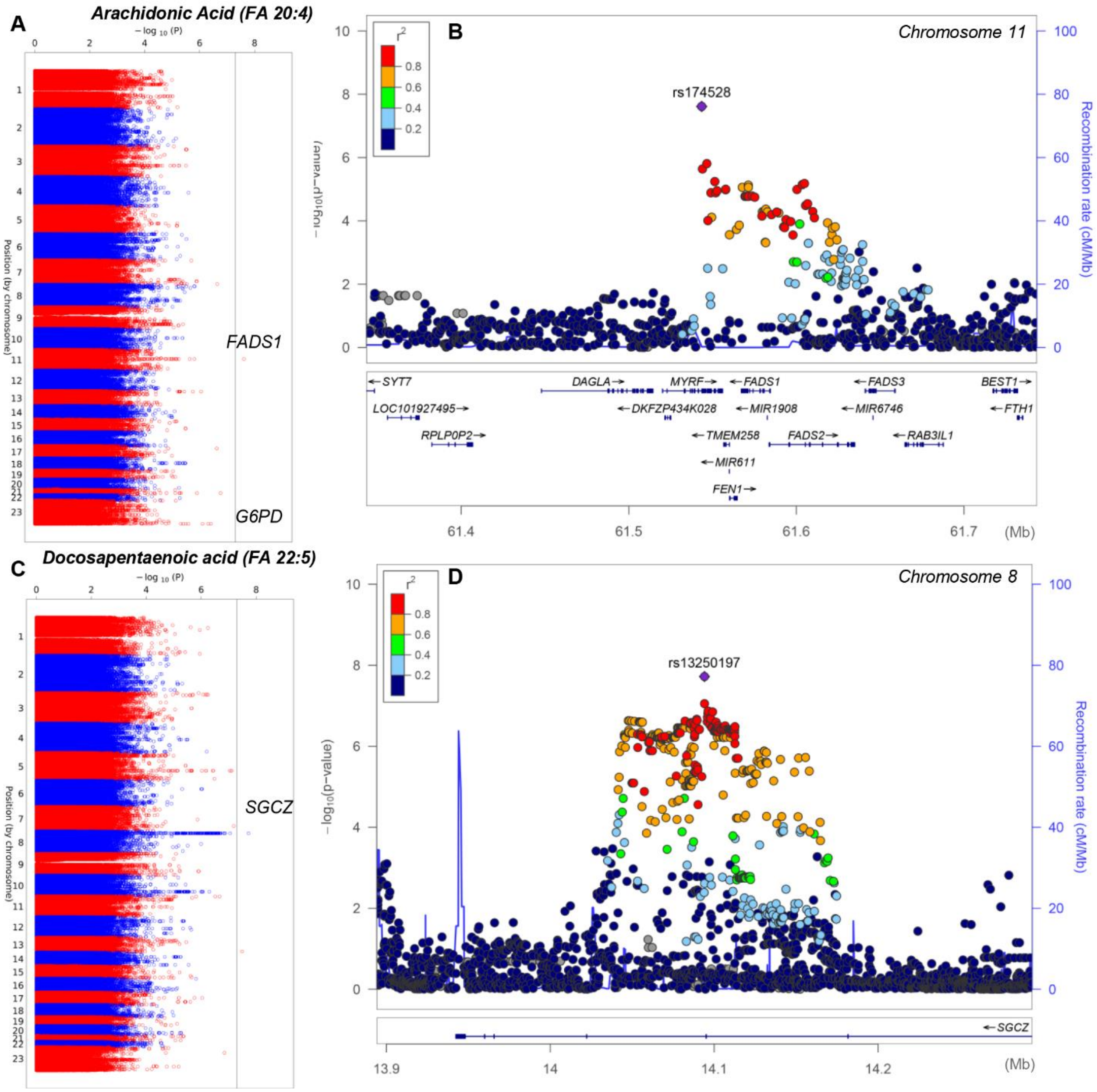


**Supplementary Figure 14 – Polymorphisms in FADS1 or G6PD and SGCZ are associated with variability in the levels of RBC highly-unsaturated fatty acids arachidonate and docosapentanoate.** Manhattan plots and LocusZooms are shown in panels **A-B** and **C-D**, respectively.

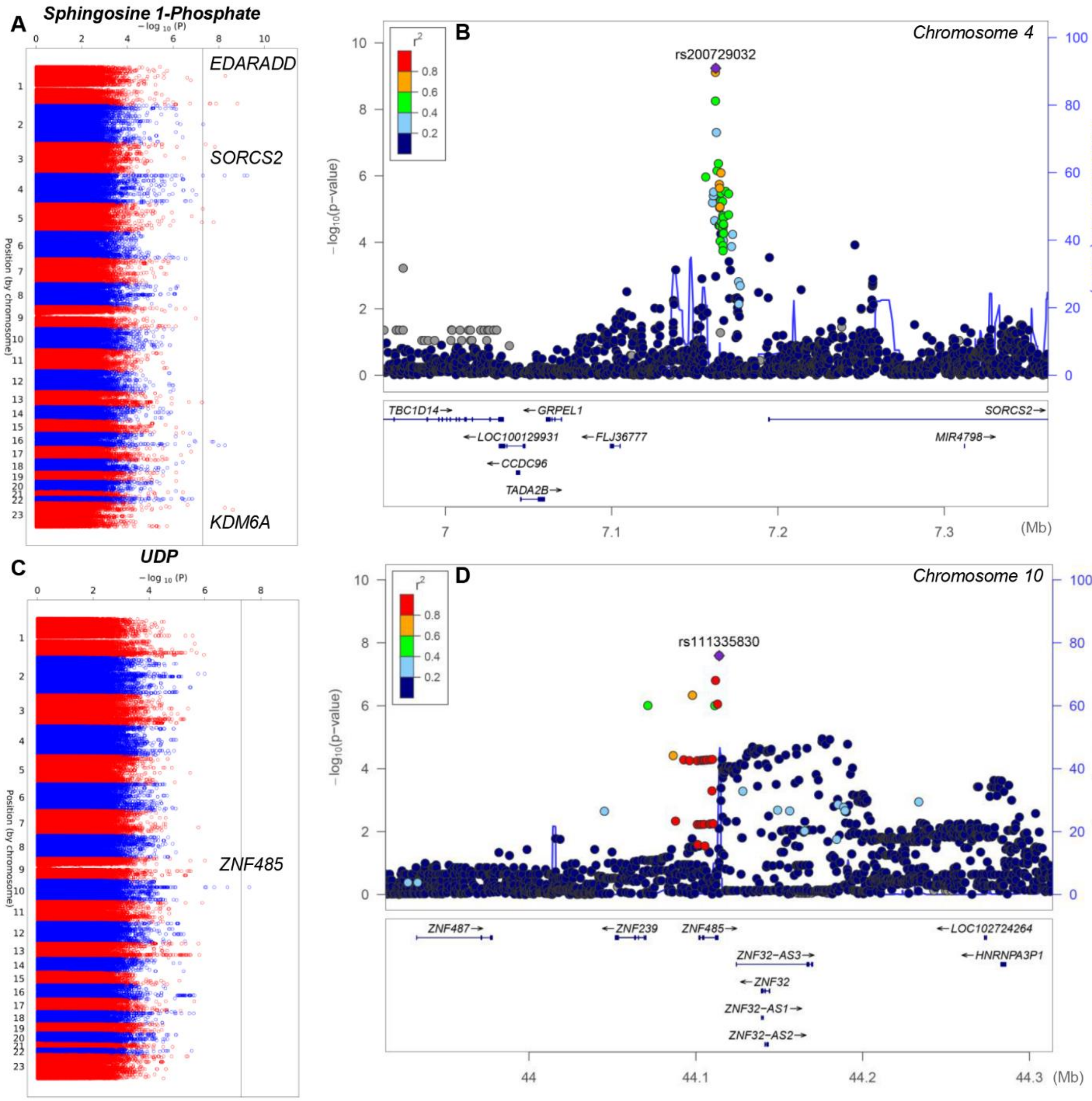


**Supplementary Figure 15 – Polymorphisms in EDARADD, SORCS2 or KDM6A and ZNF485 are associated with variability in the levels of RBC sphingosine 1-phosphate and UDP**, respectively**.** Manhattan plots and LocusZooms are shown in panels **A-B** and **C-D**, respectively.
